# Ultra-slow fMRI fluctuations in the fourth ventricle as a marker of drowsiness

**DOI:** 10.1101/2021.07.08.451677

**Authors:** Javier Gonzalez-Castillo, Isabel S. Fernandez, Daniel A. Handwerker, Peter A. Bandettini

## Abstract

Wakefulness levels modulate estimates of functional connectivity (FC), and, if unaccounted for, can become a substantial confound in resting-state fMRI. Unfortunately, wakefulness is rarely monitored due to the need for additional concurrent recordings (e.g., eye tracking, EEG). Recent work has shown that strong fluctuations around 0.05Hz, hypothesized to be CSF inflow, appear in the fourth ventricle (FV) when subjects fall asleep, and that they correlate significantly with the global signal. The analysis of these fluctuations could provide an easy way to evaluate wakefulness in fMRI-only data and improve our understanding of FC during sleep. Here we evaluate this possibility using the 7T resting-state sample from the Human Connectome Project (HCP). Our results replicate the observation that fourth ventricle ultra-slow fluctuations (∼0.05Hz) with inflow-like characteristics (decreasing in intensity for successive slices) are present in scans during which subjects did not comply with instructions to keep their eyes open (i.e., drowsy scans). This is true despite the HCP data not being optimized for the detection of inflow-like effects. In addition, time-locked BOLD fluctuations of the same frequency could be detected in large portions of grey matter with a wide range of temporal delays and contribute in significant ways to our understanding of how FC changes during sleep. First, these ultra-slow fluctuations explain half of the increase in global signal that occurs during descent into sleep. Similarly, global shifts in FC between awake and sleep states are driven by changes in this slow frequency band. Second, they can influence estimates of inter-regional FC. For example, disconnection between frontal and posterior components of the Defulat Mode Network (DMN) typically reported during sleep were only detectable after regression of these ultra-slow fluctuations. Finally, we report that the temporal evolution of the power spectrum of these ultra-slow FV fluctuations can help us reproduce sample-level sleep patterns (e.g., a substantial number of subjects descending into sleep 3 minutes following scanning onset), partially rank scans according to overall drowsiness levels, and predict individual segments of elevated drowsiness (at 60 seconds resolution) with 71% accuracy.

## Introduction

Wakefulness modulates estimates of functional connectivity, and, if unaccounted for, can become a substantial confound (Haimovici et al., 2017; Laumann et al., 2016; Tagliazucchi and Laufs, 2014). This is particularly important for clinical studies where one can expect differences in wakefulness across populations. Similarly, whether subjects keep their eyes open or closed during a given scan can also influence functional connectivity (FC) estimates (Agcaoglu et al., 2019; Dijk et al., 2010; Patriat et al., 2013). Despite such evidence, instructions regarding eye closure are not consistent across resting-state studies (Waheed et al., 2016). Moreover, eye closures (and wakefulness levels) are rarely monitored, because doing so would require some form of concurrent recordings such as eye tracking (ET; (Chang et al., 2016; Falahpour et al., 2018; Poudel et al., 2014)) or electro-encephalography (EEG; (Larson-Prior et al., 2011, 2009)). Such concurrent acquisitions are not common practice in resting-state research, and the effects of fluctuations in wakefulness (or subject’s lack of compliance with keeping eyes open) are often ignored. Prior research has demonstrated that subjects have a propensity to fall asleep during resting-state scans (Allen et al., 2014; Tagliazucchi and Laufs, 2014). It is therefore important to find ways to determine the occurrence of these events when no concurrent measures are available. This capability would not only help in future study design but may be used to assess the degree to which wakefulness fluctuations have confounded previously acquired fMRI-only datasets.

Previous work has shown that it is possible to predict sleep stages (Altmann et al., 2016; Enzo Tagliazucchi et al., 2012), and vigilance levels (Falahpour et al., 2018) relying solely on fMRI data. For example, Tagliazucchi and colleagues (2012) showed how it is possible to predict sleep stages with up to 80% prediction accuracy using dynamic traces of functional connectivity among 22 brain regions (20 cortical + 2 subthalamic nuclei) as inputs to a hierarchical tree of support linear vector machines. Similarly, significant correlation has been previously reported between fluctuations in vigilance based on EEG measures and both the global fMRI signal (Wong et al., 2013) and the average signal within a data-driven spatial template of pertinent regions (Falahpour et al., 2018). Yet, these methods have not been widely adopted by the community, either because pre-trained classifier approaches are complex (Altmann et al., 2016; E Tagliazucchi et al., 2012), or because their efficacy has only been demonstrated in small samples (e.g., 10 participants for (Falahpour et al., 2018)). It is unknown how well they may generalize beyond these small samples. Motivated by the technical need for simpler, more easily generalizable methods, and by recent work (Fultz et al., 2019) reporting that subject’s descent into sleep is accompanied by the appearance of ultra-slow (i.e., around 0.05Hz), inflow fluctuations in the fourth ventricle (*FV*), we aimed to study the ability of those fluctuations to predict wakefulness fluctuations in fMRI-only datasets, and their influence on functional connectivity. Although, as part of this evaluation, we study the spatiotemporal profile of time-locked fluctuations of equivalent frequency everywhere else in the brain, this work does not seek to settle the current debate about their etiology (i.e., neuronal or physiological). This should be the target of future enquiry, as our current results demonstrate that how one deals with these fluctuations can significantly affect inferences about how FC changes during sleep.

Our enquiry proceeded as follows. First, we study if the above-mentioned ventricular, sleep-related inflow fluctuations can be observed in an fMRI dataset (HCP dataset) not necessarily optimized to capture inflow effects. This is a key first step because the original observation of this phenomenon by Fultz and colleagues relied on data acquired at a very short TR (367ms) and with the lower boundary of the imaging field of view (FOV) sitting over the *FV*; both set this way to maximize sensitivity to inflow effects in inferior ventricular regions. This was a logical decision for their work based on the hypothesized origin of this signal—namely inflow effects due to reversal of CSF flow during sleep (Fultz et al., 2019; Grubb and Lauritzen, 2019). Yet, most fMRI data, including the HCP 7T data used here, are not acquired with such a short TR, as maximizing inflow effects is not commonly desired (Gao and Liu, 2012).

Once we confirmed the presence of these signals in the 7T HCP dataset, we evaluated if previously described relationships between those, the global signal (*GS*) and its derivative could be reproduced in this larger sample. We also searched for potential relationships with head motion and cardiac function; and studied the contribution of these ultra-slow fluctuations to the well-documented phenomena of increased global signal amplitude (*GS*_*amplitude*_) during sleep. Similarly, we also explored how and why modeling ultra-slow ventricular fluctuations as nuisance regressors might alter inferences regarding how FC changes as subject fall asleep. To finalize, we report to what degree the power density of these fluctuations can help us predict periods of drowsiness in existing resting-state samples at the sample, scan and segment levels.

Fourth ventricle ultra-slow fluctuations (∼0.05Hz) with inflow characteristics were observed during long periods of eye closure in this larger sample. The temporal evolution of these fluctuations can uncover previously reported sample-level patterns of sleep (i.e., propensity of subjects to fall asleep after 3 minutes of scanning (Tagliazucchi and Laufs, 2014), and can achieve 71% accuracy when predicting individual periods of drowsiness. Time-locked BOLD fluctuations of similar frequency were detected throughout the brain with delays in the range (min/max=-11.04/9.28 s; 5^th^/95^th^ quantile=-9.15/-2.33 s). These wide-spread BOLD fluctuations account for 50% of the increase in *GS*_*amplitude*_ that accompanies sleep. Modeling those as nuisance regressors was needed to reproduce previously reported patterns of how FC changes during eye closure and sleep (e.g., disconnection between frontal and posterior components of the DMN). In summary, this work describes how linked ultra-slow fluctuations in *FV* (inflow) and GM (BOLD) contribute to our understanding of sleep patterns in resting-state; and demonstrate their contribution in a large, commonly used fMRI sample.

## Methods

### Data

This study was conducted using a subset of the Human Connectome Project (HCP) dataset (Essen et al., 2013). We used the resting-state scans acquired on the 7T system and made publicly available as part of the 1200 Data Release (March 2018). This data subset consists of 723 different resting-state scans (15 mins each) acquired on a group of 184 subjects. This data was selected because concurrent eye pupil traces are available as part of the data release, and because the relatively short TR (1s), multi-band protocol, and high field (7T) allows for potential residual inflow effects in the *FV*. Basic scanning parameters for this data are TR=1s, TE=22.2ms, FA=45°, Voxel Resolution=1.6×1.6×1.6mm^3^, Multiband Factor=5, GRAPPA=2. Additional details can be found on the Reference Manual for the 1200 HCP Release available online at https://www.humanconnectome.org/storage/app/media/documentation/s1200/HCP_S1200_Release_Reference_Manual.pdf.

For this work, we downloaded the HCP resting-state data in two different formats:

a. The raw un-preprocessed data in original scanner space.
b. The minimally pre-processed data (Glasser et al., 2013), which includes distortion correction, motion correction and spatial normalization to the MNI template space.

In addition, we also downloaded the T1 weighted images (for visualization purposes), *Freesurfer* (Fischl, 2012) anatomical parcellations (for ROI selection), and eye tracking recordings (to be used as a proxy for fluctuations in wakefulness).

Only 404 scans from the 723 initially available were considered in this study. Table 1 lists the criteria for scan exclusion and the number of scans removed due to each criterium.

**Table 1.**
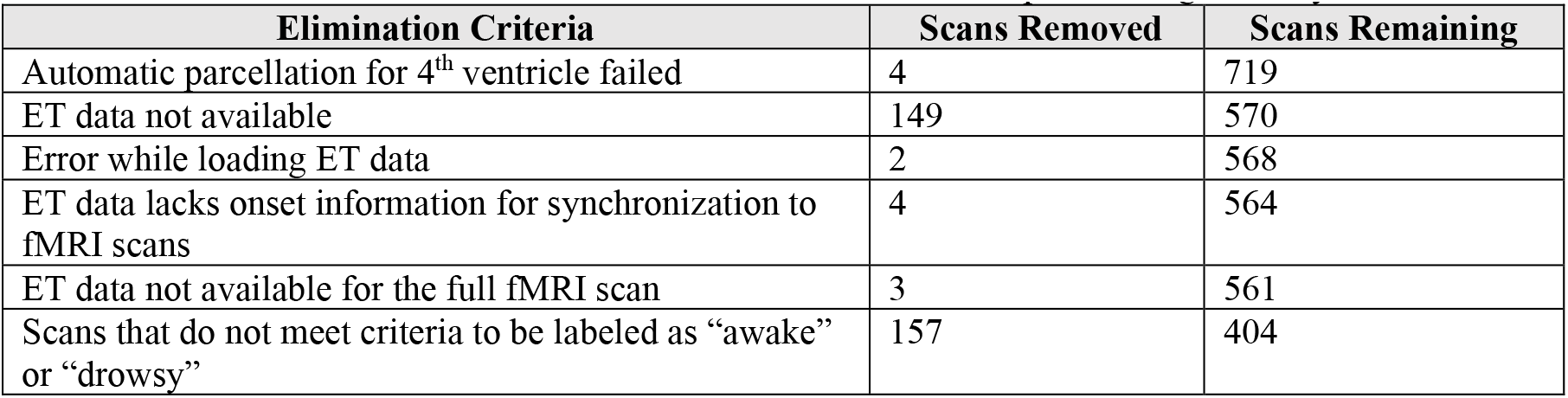
Criteria for elimination of individual scans from the final sample entering all analyses.

| Elimination Criteria | Scans Removed | Scans Remaining |
| --- | --- | --- |
| Automatic parcellation for 4<sup>th</sup> ventricle failed | 4 | 719 |
| ET data not available | 149 | 570 |
| Error while loading ET data | 2 | 568 |
| ET data lacks onset information for synchronization to fMRI scans | 4 | 564 |
| ET data not available for the full fMRI scan | 3 | 561 |
| Scans that do not meet criteria to be labeled as “awake” or “drowsy” | 157 | 404 |

### Eye Tracking

#### Eye Tracking Data Pre-processing

First, we removed pupil size samples that fell outside the temporal span of each fMRI scan to achieve temporal synchronization between the fMRI and eye tracking data. Second, we removed blink artifacts. Any period of missing pupil size data shorter than 1 second was considered a blink, and data within that period was linearly interpolated between the onset and offset of the blink. Periods of missing pupil size data longer than one second are considered eye closures and were not interpolated. Third, we observed short bursts (< 1ms) of pupil size data scattered within periods of eye closure. Those bursts were removed to ensure the continuity and correct identification of long periods of eye closure. Fourth, pupil size traces acquired at 500Hz (which is the case for 68 scans) were linearly upsampled to 1KHz to match the rest of the sample. Fifth, pupil size traces were temporally smoothed using a 200ms *Hanning* window. Sixth, pupil size traces were downsampled to 1Hz in order to match the temporal resolution of the fMRI data. Once pupil size traces were at the same temporal resolution as the fMRI data, we perform two additional operations: a) we removed the first 10 seconds of data (to match pre-processing of fMRI data outlined below) and b) we removed isolated single pupil size samples surrounded by missing data (suppl. figure 1.A), as well as interpolated isolated missing samples surrounded by data (supp. figure 1.B).

#### Scan Classification (Awake/Drowsy) based on Pupil Size Traces

We used fully pre-processed pupil size timeseries at 1Hz to classify scans in two groups:

a) <u>“*Awake*” scans</u>: defined as those for which pupil size traces indicate subjects had their eyes closed less than 5% of the scan duration.
b) *<u>“Drowsy</u>*<u>” scans</u>: defined as those for which pupil size traces indicate subjects had their eyes closed between 20% and 90% of the scan duration.

Scans with eye closure above 90% were discarded to avoid mistakenly labelling as “*drowsy*” scans that may correspond to defective eye tracking recordings (e.g., 22 scans had pupil size traces with all samples equal to zero) or to subjects that might have purposefully decided not to comply with the request to keep their eyes open from the onset of scanning.

Based on this criteria, 210 scans were labeled as “*awake*” and 194 were labeled as “*drowsy*”. The remaining scans were not used in any further analyses. Figure 1.A shows the distribution for percentage of missing eye tracking samples across the original 561 scans, and the ranges we just described for labeling of scans as “*awake*” and “*drowsy*”. Figures 1.C & D show representative pupil size traces for each type of scan. Figure 1.B shows the distributions of mean framewise displacement for both types of scans. No significant difference in head motion was found across both scan types using the *T-test* (T=-1.09, p=0.27) or the *Mann*-*Whitney* rank test (H=20412.0, p=0.97).

**Figure 1.**
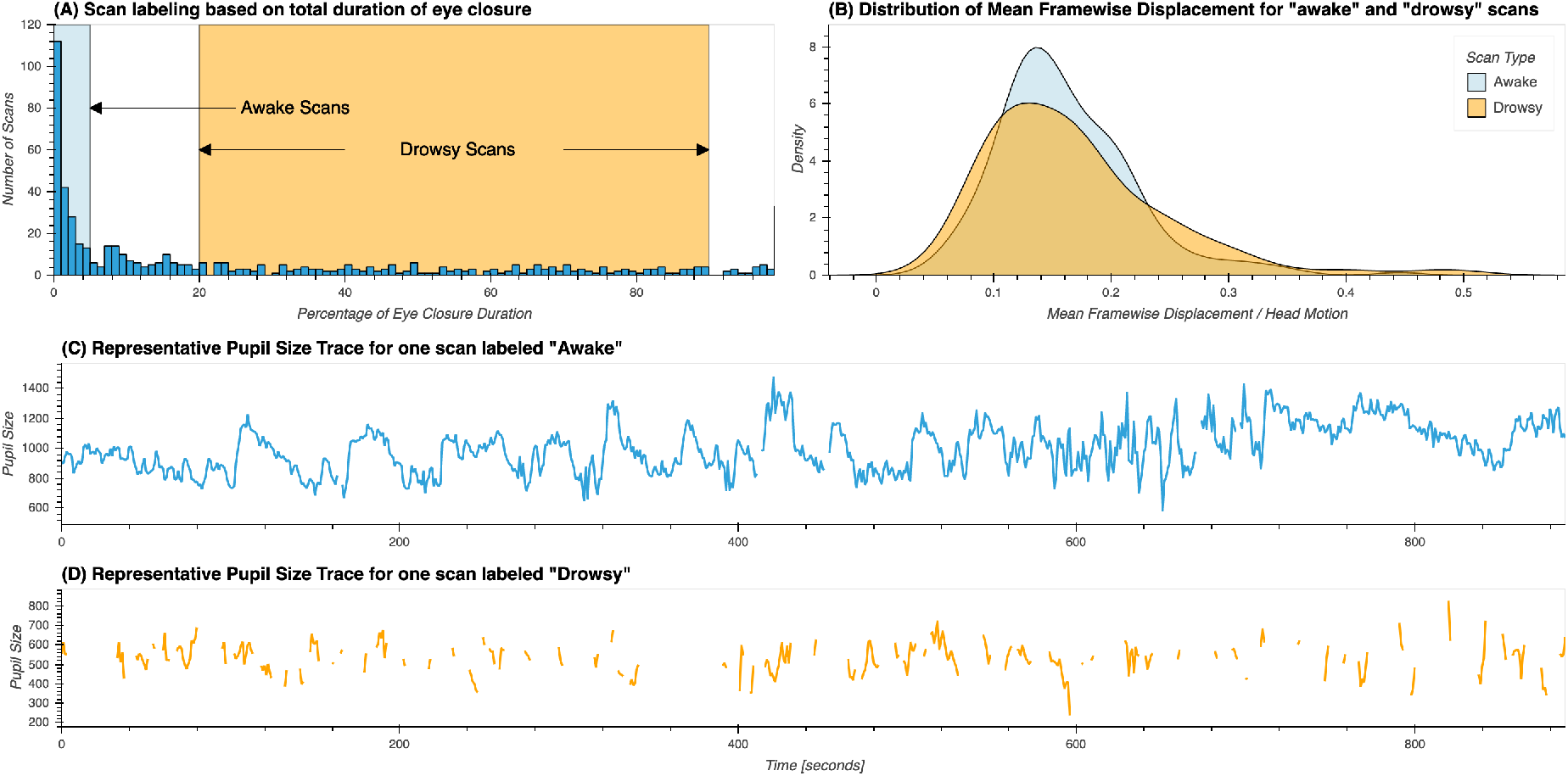
(A) Distribution of percentage of total eye closure per resting-state scan across the whole sample. Ranges used for the selection of scans marked as “*awake*” and “*drowsy*” are depicted by rectangles in light blue and orange respectively. (B) Distribution of mean values of framewise displacement across both groups of scans. (C) Presentative pupil size trace for a scan labeled as “awake”. (D) Representative pupil size trace for a scan labeled as “drowsy”.

#### Scan Segmentation based on Pupil Size Traces

For all remaining 404 scans labeled as either “*awake*” or “*drowsy*” we identified all periods of eye opening (*EO*; i.e., pupil size data available) and eye closure (*EC*; i.e., pupil size data missing); and recorded their onsets, offsets and durations. This information was used to find periods of continuous *EC* or *EO* lasting more than 60 seconds. Such scan segments were used in subsequent analyses as described below.

### Preliminary analysis on fMRI data in scanner space

Un-preprocessed fMRI data in scanner space was used for three purposes: 1) check if ultra-slow *FV* fluctuations decrease in intensity across successively collected slices; 2) evaluate confounding partial-volume effects derived from working with data transformed to MNI space later in the manuscript; and 3) estimation of cardiac traces using the “*happy*” software package (Aslan et al., 2019).

#### Across-slice signal intensity profile

Inflow fluctuations decrease in intensity across spatially contiguous slices acquired in temporal succession (Fultz et al., 2019; Gao and Liu, 2012; Yang et al., 2022). To test if such an inflow-like profile exists for signals in the *FV* in the 7T HCP dataset, we conducted some preliminary analyses on a subset of 30 scans that met the following requirements: 1) scans were labeled as drowsy based on ET traces and the procedures described above, 2) scans had an average framewise displacement below 0.1mm.

Exploration of inflow-like intensity profiles must occur in original space before any spatial transformation is applied to the data. Consequently, we cannot presume inter-scan spatial alignment beyond that derived from cautious definition of the imaging field-of-view during data acquisition. To check the level of spatial alignment among those 30 scans, we identified the location of key *FV* macro-anatomical structures in these scans. Figure 2.A shows the average of all 30 fMRI scans. Figure 2.B shows the distribution of slice numbers cutting through three key macro-anatomical structures that help define the inferior, medial and superior boundaries of the *FV*, namely the Obex (*Ob*), the posterior end of the dorsomedial recess (*pDmR*), and the inferior end of the aqueduct of Sylvius (*ASyl*). Except for one scan (black arrows in figure 2.B), there is good agreement for the location of the *FV* across the selected scans. On average, the *FV* sits between slice numbers 17 and 28 in the remaining 29 scans.

**Figure 2.**
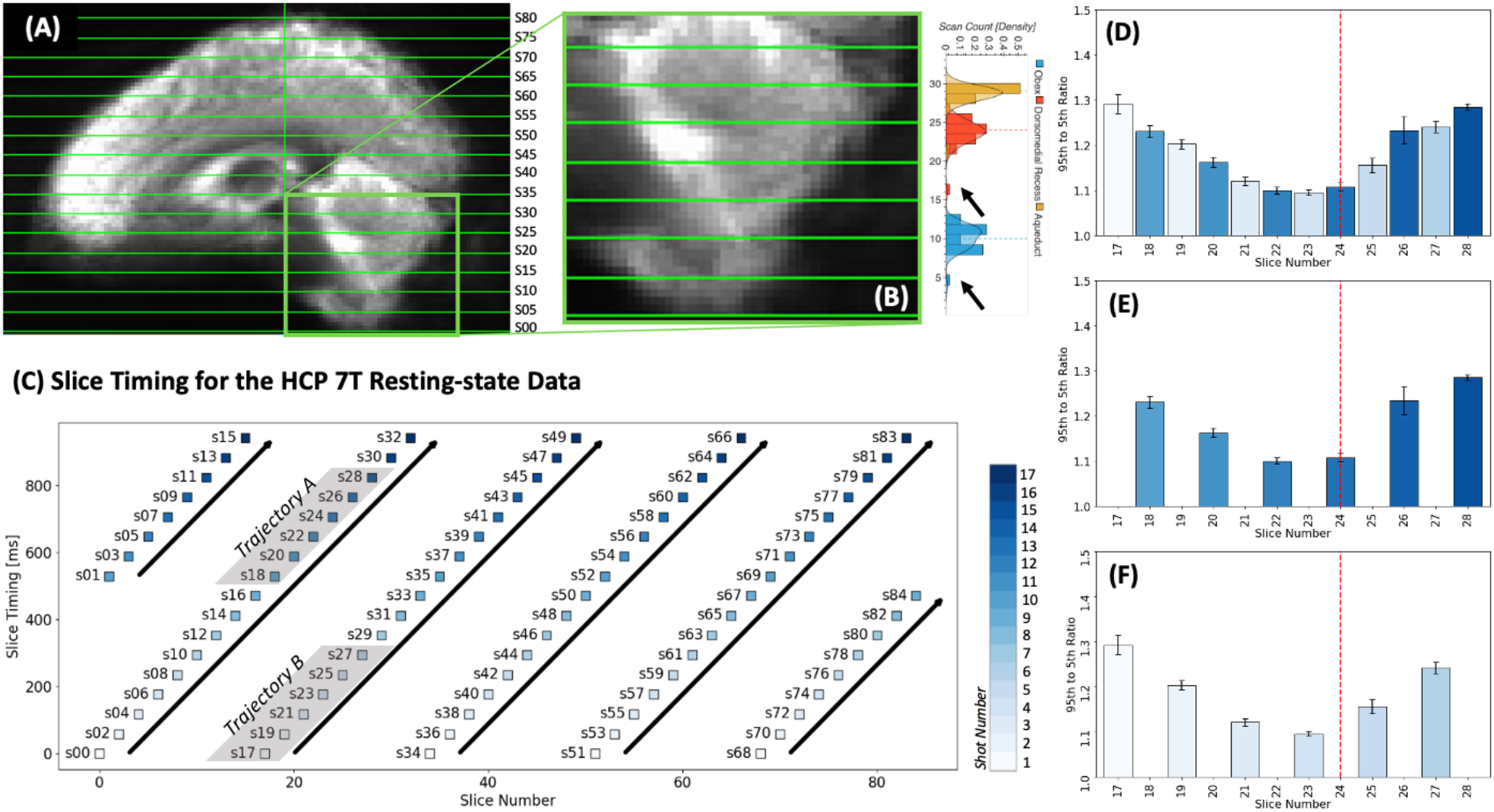
Evaluation of inflow-like properties for *FV* signals during long EC segments. (A) Average of mean timeseries is original space across the 29 scans used in these set of preliminary analyses. (B) Magnified view of the average presented in panel A. To the right we show the distribution of slice number for slices cutting through the Obex (blue), the dorsomedial recess (red) and the inferior end of the Aqueduct of Sylvius (orange). Dotted lines show the median slice number for each of these structures. Black arrows indicate the outlier location of these structures for one scan that was removed from the analyses. (C) Slice timing profile for the HCP 7T resting-state data. Each slice (labeled sXX) is represented by a colored square in a 2D plane defined by slice number/position in space (X-axis) and slice timing/position in time (Y-axis). Squares are colored according to acquisition shot. Six different ascending trajectories (tilted bold black arrows) are identified. Of these, two (grey shade) contain slices sitting over the *FV*. (D) Average 95^th^ to 5^th^ percentile *FV* signal amplitude ratio for slices crossing this anatomical structure. Bar color indicate acquisition shot number. Red dashed vertical line indicated the average location of the dorsomedial recess. (E) Same as D but only for slices in Trajectory A. (F) Same as D but only for slices in Trajectory B.

A second key consideration here is the multi-band nature of the HCP fMRI acquisitions. These data consist of 85 interleaved slices acquired using a multi-band sequence (MB Factor=5). This means that spatially contiguous slices (i.e., successive slice numbers) were not acquired consecutively on time. Figure 2.C shows a schematic of the relationship between slice number (i.e., position in space) and slice timing (i.e., position in time) for these data. In this schematic, the X-axis represents space (in terms of slice number) and the Y-axis represents time (delay in ms relative to TR onset). Each slice is represented by a square colored to reflect the shot (out of 85/5 = 17 shots) associated with it. Looking at this schematic, we learn that slices 0, 17, 34, 51 and 68 were concurrently acquired during the first shot. Similarly, we learn about six different sets of slices (or trajectories marked with black bold arrows) that were acquired consecutively and correspond to spatially increasing (although not contiguous) slices. Slices 17 through 28 (those identified in Figure 2.B as covering the *FV*) are part of two of these trajectories, labeled here as “*Trajectory A”* and “*Trajectory B”*.

Based on these considerations, we next proceeded as follows. We first manually drew *FV* ROIs in original space for each of the 29 selected scans. Next, we extracted representative time-series for each slice in these *FV* ROIs separately using the un-preprocessed data. Third, we computed (slice-wise) the ratio of the 95^th^ to the 5^th^ percentile of the signal over time during each period of eye closure longer than 60 seconds in a manner equivalent to that in Fultz et al. (2019).

Figure 2.D shows the average of these ratios across all *EC* segments for slices 17 through 28. Figure 2.E & F show the same averages but segregated into two different plots, one only including the slices in “*Trajectory A”* (2.E) and the second one those for “*Trajectory B”* (2.F). For slices in the bottom half of the *FV* (those below the *pDmR*) we observe an intensity profile compatible with inflow-like fluctuations—namely a decrease in intensity across temporally and spatially successive slices. That pattern reverses beyond that point. Given that our focus here is ultra-slow *FV* inflow fluctuations associated with sleep (i.e., those previously reported by Fultz et al. (2019)), in the rest of the manuscript we extract representative time-series for the *FV* using only voxels below the *pDmR*. We will refer to those as inferior *FV* (*iFV*) fluctuations.

#### Evaluation of partial volume effects

Because automatically generated subject-specific regions of interest for the *FV* are publicly available in *MNI* space, it is desirable to work with the data in that format. Yet, one must first ensure partial volume effects incurred during spatial normalization are not a confound.

We calculated the Pearson’s correlation between *iFV* timeseries obtained from data in *MNI* space and original space. We did this for the same set of scans used in the inflow-like profile evaluation described above. For data in original space, we manually drew *iFV* ROIs, and computed representative timeseries as the *spatial* average across ROI voxels following removal of the 10 initial acquisitions. For data in MNI space, we did the same using the group level *iFV* ROI used in the rest of the analyses (description of how this ROI was created provided below in “Inferior Fourth Ventricle Signal Extraction” section).

The average correlation value was 0.91 +/-0.06. These suggest there is strong agreement between representative timeseries obtained both ways. All other results reported in this manuscript corresponds to analyses conducted in *MNI* space.

#### Data-driven estimation of cardiac traces

The sampling frequency (*F*_*s*_) of the HCP 7T dataset is 1 Hz. The normal range for resting cardiac frequency is between 50 and 80 beats per minute (1 – 1.3 *Hz*). As such, there is potential for overlap of aliased cardiac fluctuations around 0.05 Hz (see supp. figure 2). As cardiac pulsations are known to contaminate fMRI signals recorded from ventricular compartments, it is important to evaluate their potential contribution to our observations.

The 7T HCP dataset does not include concurrent cardiac trances (e.g., ECG or pulse oximetry). We used the “*happy*” software (Aslan et al., 2019) to infer those from the raw fMRI data. Once cardiac traces were available for all 404 scans, we computed the fundamental frequency of the cardiac signal using python’s *scipy* implementation of the *Welch’s* method (*window length*=30s, *window overlap*=15s, *NFFT*=1024 samples). Next, we computed the aliased equivalent of those fundamental frequencies considering the fMRI sampling frequency of 1 *Hz*. Finally, we tested for significant differences in both original and aliased frequency profiles across scan types and segment types using both a *T-test* a non-parametric *Mann-Whitney U-test*.

### Main analyses on fMRI Data in MNI space

#### Inferior Fourth Ventricle Signal Extraction

The HCP structural pre-processing pipeline generates, among other outcomes, subject-specific anatomically based parcellations in MNI space. These parcellations are not restricted to regions within the grey matter (*GM*) ribbon, but also include subdivisions for white matter (*WM*) and cerebrospinal fluid (*CSF*) compartments, one of them being the *FV*. A group-level *iFV* region of interest (ROI) was generated by selecting voxels that are part of the *FV* in at least 98% of the sample, and then further restricting the region to only include voxels inferior to the *pDmR*. Supplementary figure 4 show this *FV* ROI overlaid on the mean EPI image across all 404 scans marked as either “*drowsy*” or “*awake*”.

To obtain scan-wise representative *iFV* timeseries, we first removed the initial 10 volumes from the minimally preprocessed data, and then scaled the data by dividing each voxel timeseries by its own mean and multiplying by 100. The representative *iFV* timeseries for each scan was computed as the average across all voxels in this ROI. The representative timeseries for this region was extracted prior to any additional pre-processing to minimize the removal of inflow effects and also minimize partial volume effects.

#### Frequency Analysis of the iFV Signal

First, we estimated the power spectral density (*PSD*) for each subject’s *iFV* timeseries using python’s *scipy* implementation of the *Welch’s* method (*window length*=60s, *window overlap*=45s, *NFFT*=128 samples). We did this both at the scan level (i.e., using complete scan timeseries) and the segment level (i.e., using only segments of continuous *EO* and *EC*, as described above). We then compared *PSD* across scan types (“*awake*” vs. “*drowsy*”) and segment types (*EC* vs. *EO*). Significant differences were identified using the *Kruskal-Wallis H-test* at each available frequency step and *p*_*Bonf*_ <0.05, corrected for the number of different frequencies being compared.

Next, to study the temporal evolution of the *iFV* signal fluctuations centered around 0.05Hz, we also computed spectrograms (i.e., *PSD* across time) using python’s *scipy spectrogram* function (*window length*=60s, *window overlap*=59s, *NFFT*=128). We did so for each scan separately, and then computed the average temporal traces of PSD within two different frequency bands:

a) *<u>“Sleep</u>*<u>” band</u>: 0.03 – 0.07 Hz.
b) *<u>“Control</u>*<u>” band</u>: 0.1 – 0.2 Hz.

The range of the sleep band was selected based on the original work of Fultz et al. (2019) so that it would include the target frequency of interest (0.05Hz). The control band was selected as to exclude respiratory signals as well as the 0.01 – 0.1 Hz band, which contains the majority of neuronally-induced fluctuations in resting-state data (Cordes et al., 2001). In later analyses we refer to the total area under the *PSD* trace for the sleep band as *PSD*_*sleep*_ and for the control band as *PSD*_*control*_.

Supplementary figure 5 depicts this process. Panel A shows the average BOLD signal in *iFV* for one resting-state scan from a representative subject. Panel B shows the associated spectrogram, as well as the sleep (green) and control (red) bands. Finally, panel C shows the average *PSD* within these two bands of interest (green and red curves). For this scan, a strong fluctuation around 0.05Hz appears during the second half of the scan (panel A). This fluctuation results in an increased *PSD* for a small range of frequencies around 0.05Hz (panel B). Such increase becomes clearly apparent when we look at the average *PSD* traces for the “*sleep*” and “*control*” bands in panel C. While the green trace (“*sleep*” band) shows large positive deflections during the second half of the scan, that is not the case for the red trace (“*control*” band).

#### Additional fMRI pre-processing

The HCP minimal preprocessing pipeline (Glasser et al., 2013) includes steps to account for distortion correction, motion correction and spatial normalization to the MNI template space, yet it does not include other important common pre-processing steps (e.g., spatial smoothing, temporal filtering, nuisance regression). The following additional pre-processing pipelines were used in this work:

##### Smoothing Pipeline

minimally pre-processed pipeline followed by 1) discarding of initial 10 volumes, 2) spatial smoothing (*FWHM=4mm*), 3) intensity normalization dividing by the voxel-wise mean timeseries, 4) regression of slow signal drifts (Legendre polynomials up to the 5th order), and 5) band-pass filtering (0.01 – 0.1 *Hz*).

##### Basic Pipeline

same as the *Smoothing* pipeline, but the regression step also includes motion parameters and their first derivative as additional regressors.

##### CompCor Pipeline

same as the *Basic* pipeline, except that a set of additional regressors based on the *CompCor* technique (Behzadi et al., 2007) were included to model physiological noise. Those regressors are the first five principal components of the signal in the ventricles and WM (see supplementary figure 6). Subject specific masks for these regions were created by combining the respective subject-specific masks for these compartments generated by *Freesurfer* (Fischl, 2012), and then eroding those by one voxel.

For all three pipelines, we also worked with a modified version (i.e., the *“+”* version) that includes an additional voxel-wise regressor corresponding to a shifted version of the *iFV* signal with maximal correlation in each voxel. We refer to those pipelines as the ***Smoothing+, Basic+*** and ***CompCor+*** pipelines in the remaining of the text.

All pre-processing steps, beyond those part of the minimally pre-processing pipeline, were performed with the AFNI software (Cox, 1996).The regression step in all pipelines was always performed at the whole-scan level, not on a segment-by-segment basis.

#### Voxel-wise correlation analyses: zero-lag

Zero-lag correlation analyses were based of the *Basic* pipeline. Voxel-wise maps of correlation between the *iFV* signal (filtered to the same range as the rest of the data [0.1 to 0.01 *Hz*]) and all voxels in the full brain mask (available as part of the HCP data release) were computed separately for each period of *EC* and *EO* longer than 60 seconds. Next, we generated average correlation maps per eye condition by averaging the individual maps for all segments of a given type. For this, we applied a Fisher *Z* transform prior to averaging. Following averaging, Z-values were converted back to R-values prior to presentation. We also performed a *T-test* across the population (*AFNI* program *3dttest*+) to identify regions with significant differences in correlation with the *iFV* signal across eye conditions. Finally, histograms of correlation values per eye condition were computed based on the average maps.

#### Voxel-wise correlation analyses: cross-correlation and lag maps

The *RapidTide2* software (https://github.com/bbfrederick/rapidtide ; v2.0.8+11.gb17c48f) was used to perform cross-correlation analyses. Given, the *Rapidtide2* software applies bandpass filtering (also set to 0.01 – 0.1 *Hz*), inputs to this software were the data from the *Basic* pipeline, excluding the filtering step.

In addition to voxel-wise cross-correlation traces, the *Rapidtide2* software also produces a) voxel-wise lag maps, where each voxel is assigned the temporal lag leading to the strongest positive correlation between the source signal (i.e., that from the *iFV*) and the signal in that particular voxel; b) masks of statistical significance for cross correlation results based on non-parametric simulations with null distributions (n=10,000), c) and time-shifted voxel-wise versions of the source signal leading to maximal correlation.

Cross-correlation analyses were conducted at the scan level using only scans labeled as “*drowsy*”, as those are expected to contain more fluctuations of interest (i.e., those associated with light sleep). To generate group-level lag maps and voxel-wise cross-correlation traces we computed the average across scans taking into account only results rendered significant at *p<0*.*05* at the scan-level. We only show results for voxels where at least 40% of the scans produced significant cross-correlation results. This averaging approach was followed to avoid non-significant low correlations and noisy lag estimation contributing to the group level results.

#### Relationship to the Global Signal

In fMRI, *GS* refers to the spatial average of voxel-wise timeseries across the brain. GS is a combination of multiple signal sources, and its nature is not fully understood (see (Liu et al., 2017) for an excellent review on the topic). One often reported finding is that its amplitude (*GS*_*amplitude*_) significantly increases during sleep compared to awake states (Fultz et al., 2019; Horovitz et al., 2008; Larson-Prior et al., 2009). Similarly, an inverse relationship is expected between *FV* inflow effects and the derivative of the *GS* if we assume that CSF dynamics are driven by changes in cerebral blood volume (Fultz et al., 2019; Yang et al., 2022).

We investigated if we could observe these effects on our sample. First, we computed the *GS* as the average signal across all grey matter ribbon voxels (as in (Fultz et al., 2019)). Second, we computed the negative zero-thresholded time derivative of the *GS* as the derivative of the *GS* multiplied by −1 and with all negative values set to zero to restrict the analysis to inflow, not outflow. This is the same procedure used by Fultz et al. (2019). We refer to this signal as *-dGS/dt* in the rest of the manuscript. Third, we conducted cross-correlation analysis (lags in the range [ - 20, 20 s]) to see if we could reproduce previously reported temporal lag relationships between these three signals (i.e., GS, -dGS/dt and *iFV*).

Finally, to explore the contribution of GM ultra-slow fluctuations time-locked to those in *iFV* to *GS*_*amplitude*_ and its increase during sleep, we estimated segment-level *GS*_*amplitude*_ in terms of the temporal standard deviation. Significant increases in *GS*_*amplitude*_ between *EO* and *EC* segments were evaluated via *T-test*.

#### Effects on inter-regional FC

To study how removal of *FV* fluctuations may affect inferences about changes in FC during sleep, we computed whole-brain FC matrices using the 200 ROI Schaefer Atlas (Schaefer et al., 2017) for three pre-processing pipelines: *Basic, CompCor* and *CompCor*+.

For each ROI, a representative timeseries was generated as the average across all voxels part of that ROI. Statistical differences in connectivity between “*drowsy*” and “*awake*” scans were then evaluated using Network-based Statistics (Zalesky et al., 2010) using T=3.1 as the threshold at the single connection level, and *p<0*.*05* at the component level (5000 permutations). Finally, results are presented in brain space using the *BrainNet Viewer* software (Xia et al., 2013).

#### Classification Analyses

To study how well the presence of fluctuations around 0.05Hz in the *iFV* can help us predict reduced wakefulness, we used a logistic regression classifier that takes as inputs windowed (window length = 60s, window step = 1s) estimates of *PSD*_*sleep*_. A duration of 60s was selected to match the frequency analyses described before. Classification labels were generated as follows. For each sliding window, we calculated the percentage of eye closing time for that window. If that percentage was above 60%, we assigned the label “*eyes closed/drowsy*”. If the percentage was lower than 40%, we assigned the label “*eyes open/awake*”. All other segments were discarded given their mix/uncertain nature. The input to the classifier was the average *PSD*_*sleep*_ in the corresponding window.

This analysis was conducted using all 404 scans. This procedure resulted in an imbalanced dataset with 311184 samples, of which 233451 (∼75%) correspond to “*eyes open/awake*” class, and the rest (77733; ∼25%) to the “*eyes closed/drowsy*” class. To account for this imbalance, we used a stratified k-fold approach to estimate the generalizable accuracy of the classifier (k=10) and assigned class weights inversely proportional to the frequency of each label during the training. Those analyses were conducted in Python using the scikit-learn library (Abraham et al., 2014).

The same analysis was conducted also using as input the sliding window version of the *GS*_*amplitude*_, a randomized version of the *PSD*_*sleep*_ (control case) and using as features both the windowed *GS*_*amplitude*_ and the *PSD*_*sleep*_. Significant differences in accuracy across these scenarios was evaluated via T-test (with as many accuracy estimates per classification scenario as k-folds). Additionally, a representative confusion matrix per classification scenario was created by computing median precision values across k-folds.

### Data and Code availability statement

The data used in this work is a subset of the Human Connectome Project (Essen et al., 2013) 7T resting-state sample that is publicly available. Analyses were conducted with publicly available packages including *AFNI, RapidTide, Python*, and *Scikit-learn*. Processing scripts publicly available at https://github.com/nimh-sfim/hcp7t_fv_sleep

## Results

### Eye Tracking

Figure 3 shows the percentage of scans (Y-axis) for which subjects had their eyes closed at a particular fMRI volume acquisition (X-axis). Individual dots represent actual scan counts at each TR, while the black line represents a linear fit to the data. For approximately 10% of scans, subjects had their eyes closed during the first fMRI acquisition. As scanning progresses, the number of scans for which subjects had their eyes closed increases (Linear Fit R=0.96). By the end of the resting state scans, in approximately 40% of scans, subjects had their eyes closed; signaling that as time inside the scanner advances more subjects may have fallen asleep.

**Figure 3.**
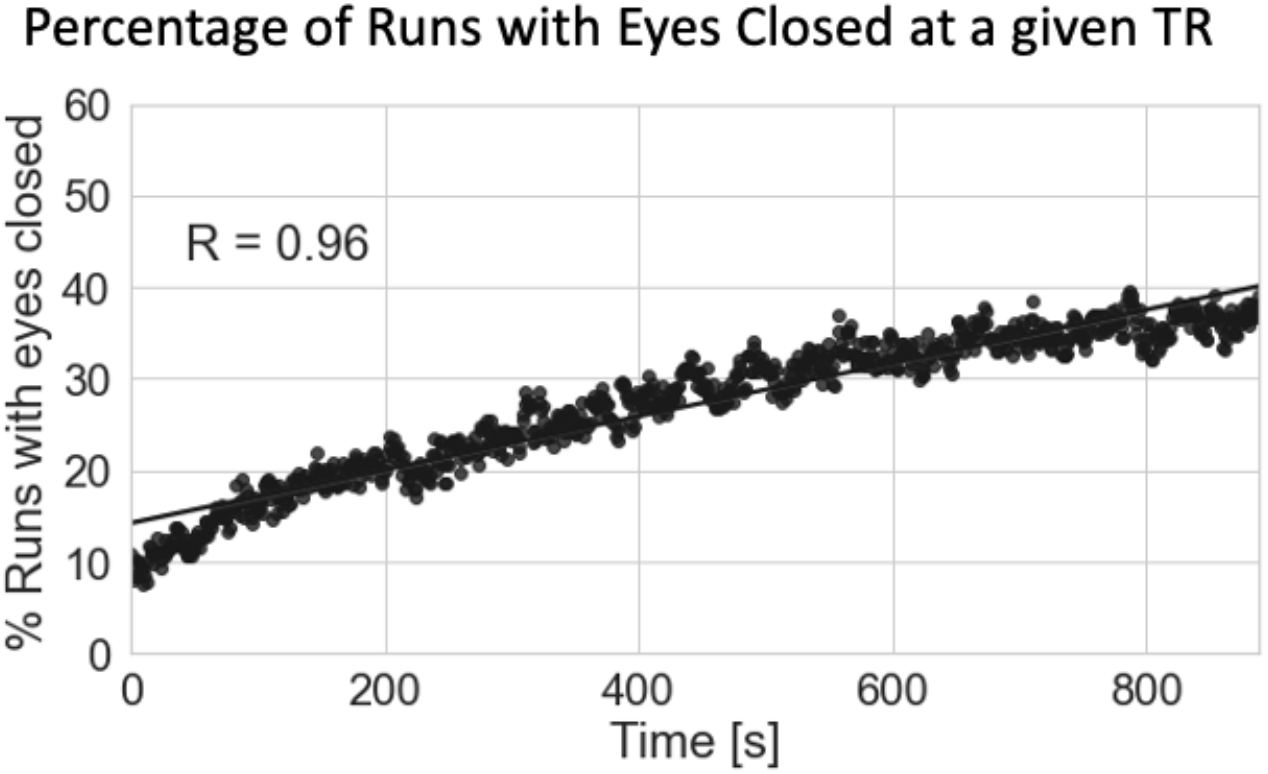
Percentage of scans with eyes closed at a given fMRI acquisition. As scanning progresses, the number of subjects with their eyes closed at a given fMRI acquisition (i.e., TR) increases.

### Relationship to Cardiac Function

Figure 4.A shows the distribution of scan-level heart rates for all 404 scans. For most scans, heart rate falls within the normal ranges of resting heart rate (red area), suggesting the “*happy*” software extracted valid cardiac traces. Figure 4.B shows the distribution of aliased cardiac rate given fMRI’s sampling frequency of *1Hz*. As predicted by the simulations (supplementary figure 2), for some scans, aliased cardiac frequency falls within our target frequency range of interest (0.03 Hz to 0.07Hz; green area). Figure 4.C & D shows box plots of cardiac frequency and aliased cardiac frequency by scan type respectively. While a significant difference was observed in cardiac frequency between awake and drowsy scans (*T=2*.*24, p=0*.*03* | *U=23162, p=0*.*02*), no such difference was found for their aliased equivalents (*T=1*.*27, p=0*.*21* | *U=21507, p=0*.*33*).

**Figure 4.**
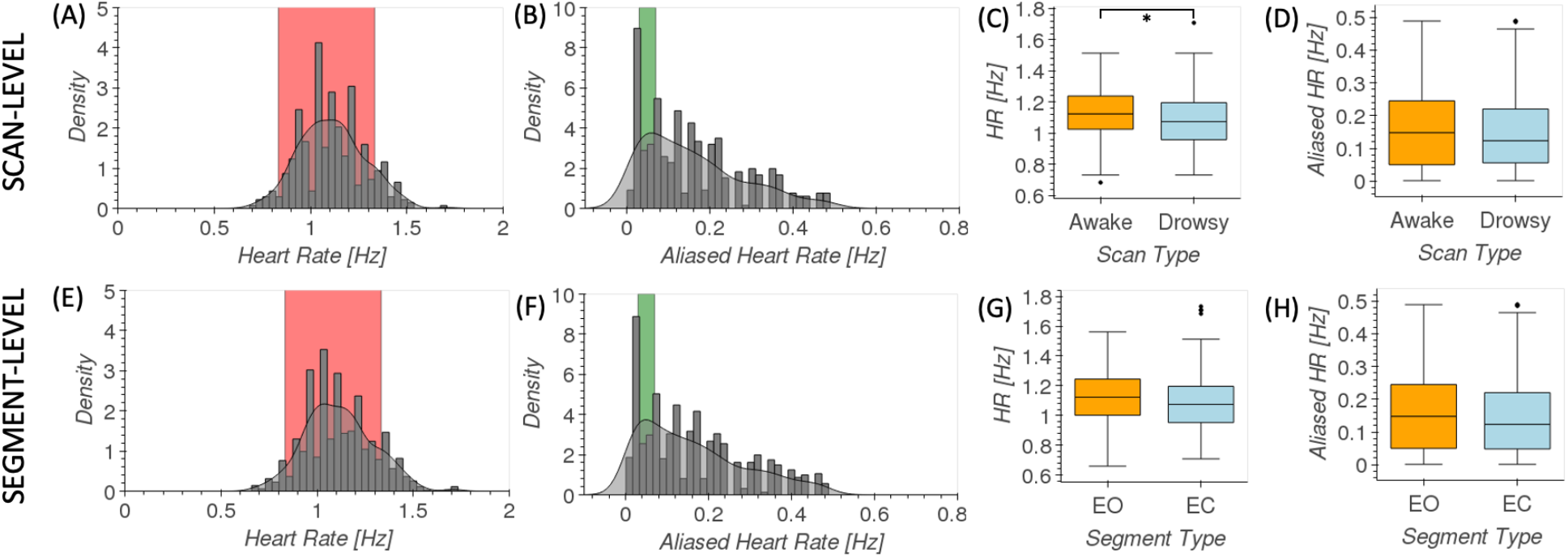
Analysis of data-driven estimates of cardiac traces. (A) Distribution of scan-level heart rate across all scans. Red indicates common ranges for resting heart rate. (B) Distribution of aliased scan-level heart rate across all scans. Green indicates the target frequency range in fMRI recordings for sleep-related fluctuations in the *FV*. (C) Box plots of scan-level heart rates for awake (orange) and drowsy (light blue) scans. (D) Box plots [middle line = median, box edges = 25-75 quantiles, whiskers = furthest data point within 1.5 times the interquartile range above and below the box edge] of scan-level aliased heart rates for awake (orange) and drowsy (light blue) scans. (E) Distribution of segment-level heart rate across all scans. Red indicates common ranges for resting heart rate. (F) Distribution of aliased segment-level heart rate across all scans. Green indicates the target frequency range in fMRI recordings for sleep-related fluctuations in the *FV*. (F) Box plots of segment-level heart rates for EO (orange) and EC (light blue) segments. (D) Box plots of segment-level aliased heart rates for EO (orange) and EC (light blue) segments.

Similarly, figures 4.E-H show equivalent information at the segment level. Again, a significant difference for cardiac rate between long segments of EO and EC (*T=2*.*34, p=0*.*02* | *U=100228, p=0*.*002*) was observed, but not in terms of their aliased equivalents (*T=1*.*71, p=0*.*09* | *U=95061, p=0*.*08*).

### Relationship to Global Signal

Figure 5.A shows the distribution of *GS*_*amplitude*_ for *EC* and *EO* segments for all pre-processing pipelines, and that of *iFV* amplitude. In all instances, amplitudes are significantly larger during *EC* segments compared to *EO* segments according to *T-tests* and p_*Bonf*_ <0.05 (see supp. table 1 for details). Yet, in absolute terms, differences in *GS*_*amplitude*_ went down in half from the *Basic* pipeline (0.15) to the *CompCor*+ pipeline (0.07).

**Figure 5.**
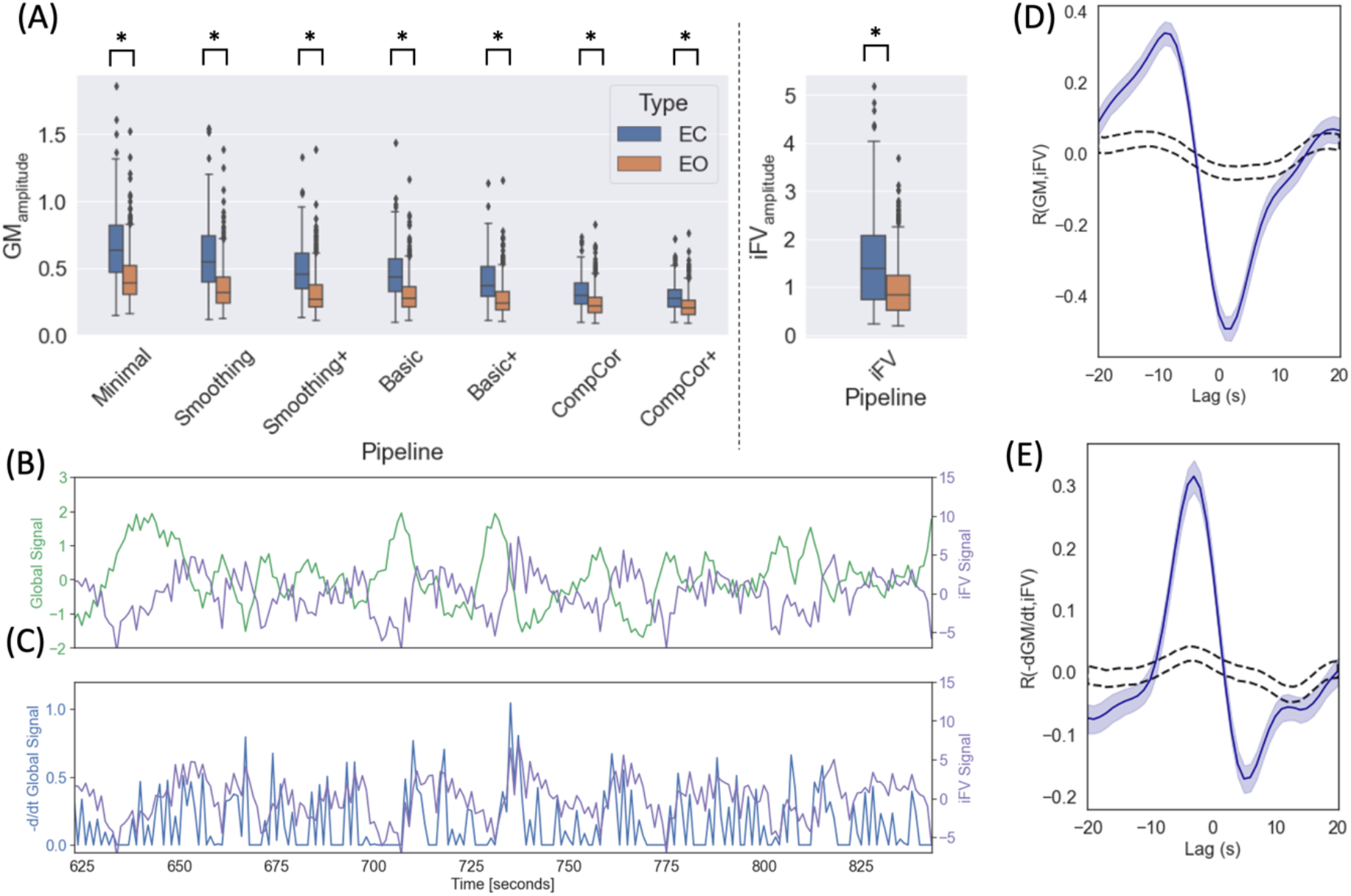
Relationship between *GS, -d/dtGS* and *iFV* signal. (A) Segment-level *GS*_*amplitude*_ for different pre-processing pipelines and for the *iFV* signal. In all instances we observed a significant difference across segment types. (B) *iFV* and GS signal for one representative *EC* segment. (C) *iFV* and *-d/dtGS* for the same representative EC segment. (D) Cross-correlation between *GS* and *iFV* signal (blue line) (E) Cross-correlation between *-d/dtGS* and *iFV* signal (blue line). In both (D) and (E) shaded blue regions indicate 95% confidence interval for the actual data and dashed black lines indicate 95% confidence interval when data is shuffled (n=1000 permutations).

Figure 5.B-E shows how we can reproduce prior observations regarding the relationship between *GS, -dGS/dt* and *iFV* made by Fultz and colleagues in our sample. Figure 5.B shows, for a representative EC segment, both the *GS* (green) and the *iFV* signal (magenta). These two signals appear to be anticorrelated. Next, figure 5.C shows, for the same representative *EC* segment, both the *-dGS/dt* (blue) and the *iFV* signal (magenta). In this case, both signals appear to be positively correlated. Figure 5.D shows the average cross-correlation profile between the *GS* and *iFV* signals [*Max. corr* = −0.49 | *lag* = 2s] across all EC segments. Figure 5.E shows the average cross-correlation profile between *-dGS/dt* and *iFV* signals [*Max. corr*=0.32, lag = −3s] across all EC segments.

### Spectral Characteristics of the iFV signal

#### Power Spectral Density (PSD) of the iFV signal

Figure 6.A shows average PSD across all scans labeled as “*awake*” (orange) and those labeled as “*drowsy*” (blue). Figure 6.B shows average PSD across all scan segments during which subjects had their eyes closed (EC; blue) for at least 60 seconds and for all scan segments during which subjects had their eyes open (EO; orange) for the same minimum duration. We can observe that PSD is significantly higher for “*drowsy*” scans as compared to “*awake*” scans. The same is true for EC segments as compared to EO segments (*p*_*Bonf*_ *<0*.*05*). Significant differences concentrate primarily below 0.06Hz at the scan level, and extents all the way to 0.08Hz when looking at the segment level.

**Figure 6.**
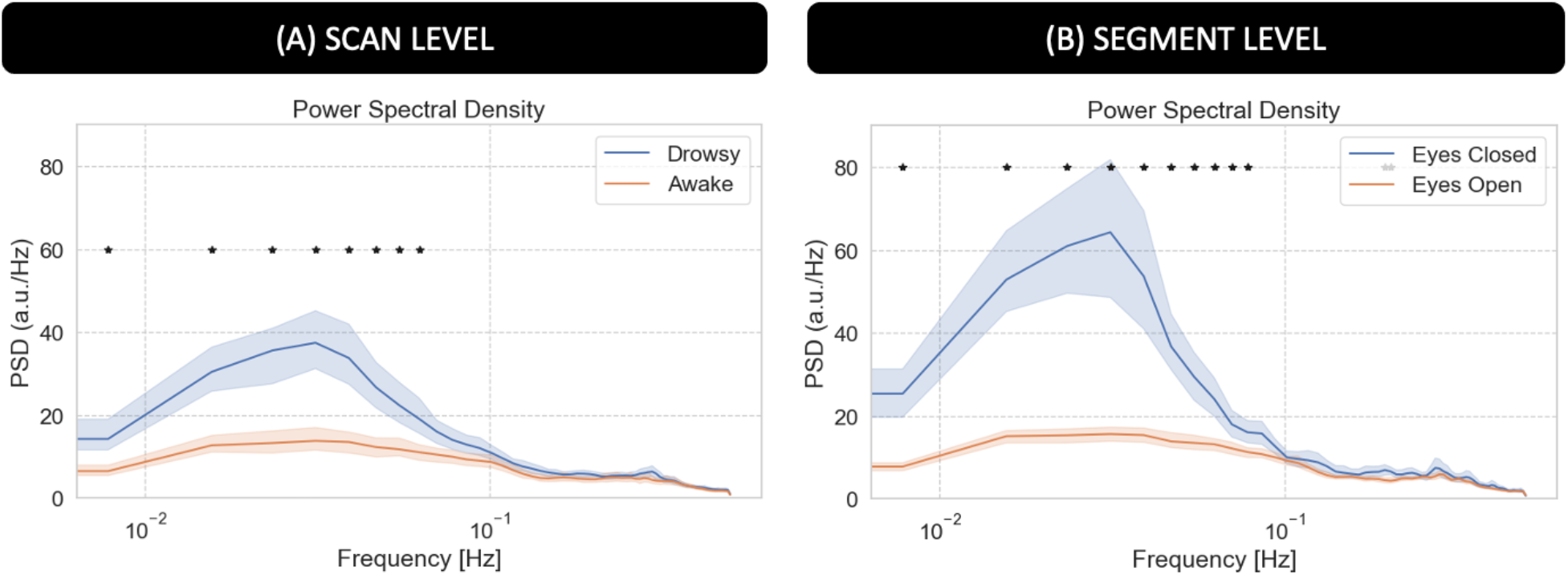
Power Spectral Density (PSD) analysis for the average fMRI signal in the 4^th^ ventricle. (A) Average PSD for “awake” and “drowsy” scans. (B) Average PSD across eyes closed and eyes open segments. In both panels, frequencies for which there is a significant difference (p_Bonf_ <0.05) are marked with an asterisk.

#### Time-frequency analyses for the iFV signal

Figure 7 summarizes our results from the time-frequency analyses for the *iFV* signal. First, figure 7.A shows the temporal evolution of the average Power Spectral Density (*PSD*) in two different bands—namely the sleep band (*PSD*_*sleep*_) and control band (*PSD*_*control*_)—for both “*awake*” and “*drowsy*” scans. *PSD*_*sleep*_ evolves differently with time for both scan types. For “*awake*” scans, *PSD*_*sleep*_ remains relatively flat for the whole scan duration. Conversely, for “*drowsy*” scans, the *PSD*_*sleep*_ shows an incremental increase as scanning progresses. This increase becomes clearly apparent after the initial 3 minutes of scan. No difference in the temporal evolution of the *PSD*_*control*_ was observed across scan types.

**Figure 7.**
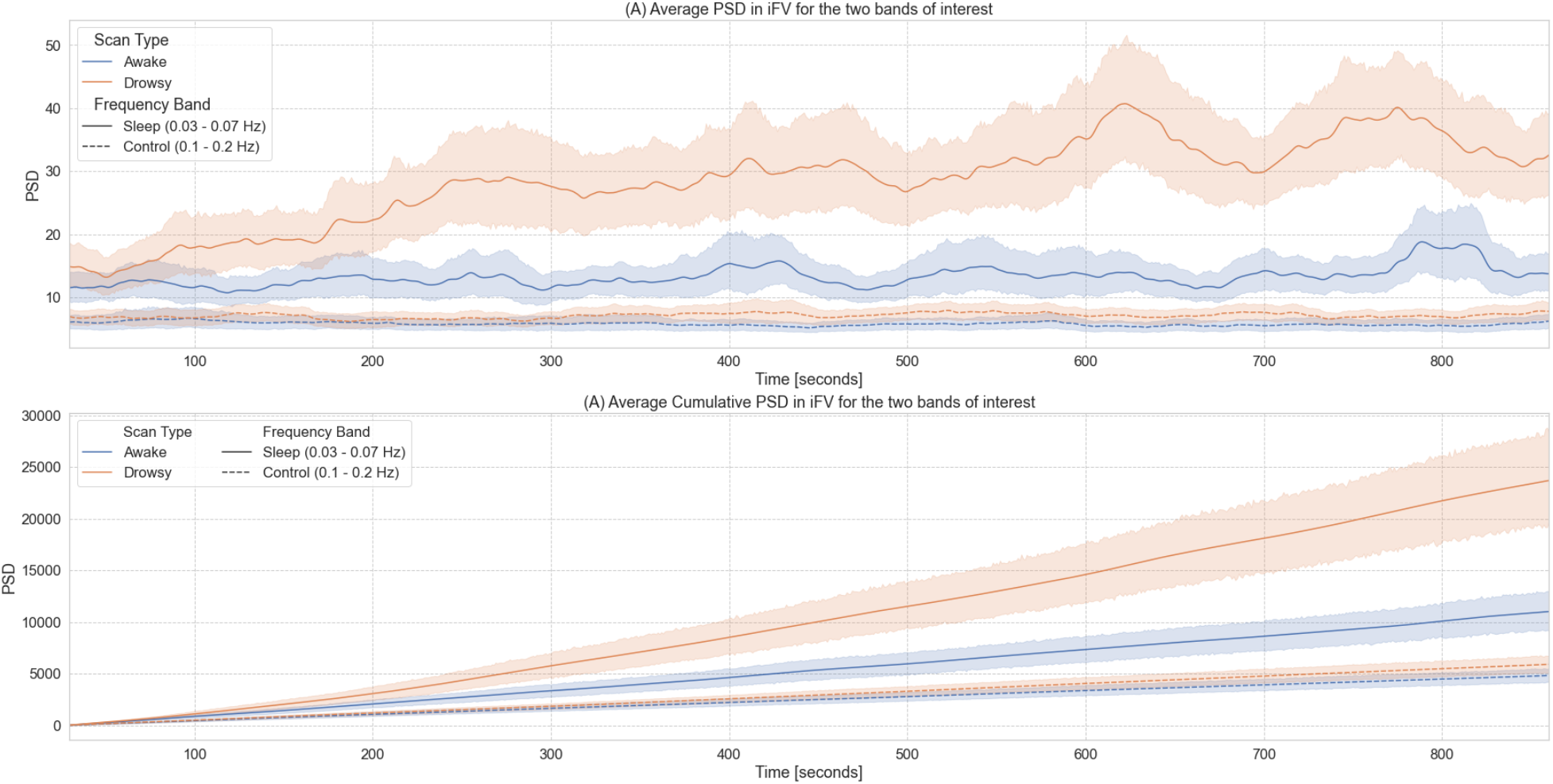
Time-frequency analyses for the BOLD signal from the 4^th^ ventricle. (A) Temporal evolution of the power spectral density (PSD) of this signal in two different frequency bands (sleep band: 0.03 – 0.07Hz, control band: 0.1 – 0.2Hz) for scans labeled as “awake” and “drowsy”. (B) Cumulative PSD for the same two bands and scan types shown in the top panel. In both panels, lines show the average across all scans of a given type, and shaded regions indicate 95% confidence intervals (bootstrapping n=1000). Continuous lines correspond to trends for the sleep band, while dashed lines corresponds to the control band.

As subjects are not expected to close their eyes or fall slept in synchrony, we also looked at cumulative versions of *PSD*_*sleep*_ and *PSD*_*control*_ as a function of time (Figure 7.B). In this case, we observe a faster increase in cumulative *PSD*_*sleep*_ for “*drowsy*” scans as compared to “*awake*” scans. This difference becomes clearly apparent approximately after 180 seconds. Conversely, for the cumulative *PSD*_*control*_, we did not observe any difference in the rate of increase across scan types.

### Voxel-wise zero-lag correlations

For periods of eye closure (EC, Figure 8.A), we observe wide-spread negative correlations between the sleep-related signal in the *iFV* and grey matter (colder colors). We also observe positive correlation in periventricular regions superior to the *FV* (warmer colors). Importantly, not all regions correlate with the same intensity. The strongest negative correlations (r < −0.3) concentrate on primary somatosensory regions, primary auditory cortex, inferior occipital visual regions and the occipito-parietal junction. The strongest positive correlations (r > 0.3) appear mostly on the edges of ventricular regions above *FV*.

**Figure 8.**
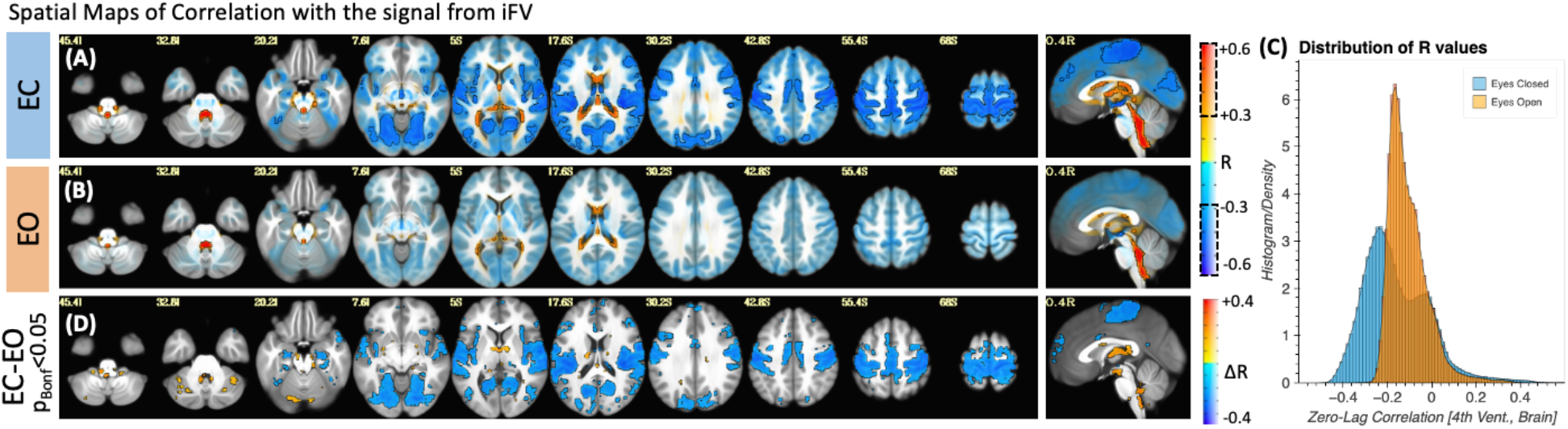
(A) Group-level map of voxel-wise average correlation between the iFV signal and the rest of the brain for periods of eye closure longer than 60 seconds. (B) Same as (A) but for periods of eye opening longer than 60 seconds. In both (A) and (B), areas with | R | > 0.3 are highlighted with a black border and no transparency. Areas with |R| < 0.3 are shown with a higher level of transparency. (C) Histogram of voxel-wise average R values across all scan segments for both conditions. A shift towards stronger correlation values can be observed for the eyes closed condition. (D) Significant differences in voxel-wise correlation between eyes closed and eyes open condition (p_Bonf_ <0.05).

For eye opening (*EO*) periods (Figure 8.B), although there is a mostly negative relationship between the signal from the 4^th^ ventricle and the rest of the brain, this relationship is weaker (|r|<0.3 mostly everywhere). Figure 8.C shows the distribution of correlation values for both *EC* (blue) and *EO* (orange). While for *EO* correlation values are mostly centered around zero; for *EC* we can see a bimodal distribution with one peak around −0.3 (corresponding the areas of strong negative correlation within grey matter) and a second positive peak around 0.05 with a long positive tail (corresponding to the strong positive correlations in other ventricular regions). Finally, a test for statistical difference (*p*_*Bonf*_ *<0*.*05*) in voxel-wise correlation values between the *EO* and *EC* segments (Figure 8.D) revealed that regions with significant differences mimic those shown in figure 8.A as having the strongest correlations during *EC*.

### Voxel-wise temporal delays

To explore differences in arrival time across the brain, we also conducted voxel-wise cross correlation analyses based on the *iFV* signal using only the “*drowsy*” scans. Figure 9.A shows the resulting average map of temporal lags, where the color of a voxel represents the lag conductive to the maximum positive cross-correlation value at that location. Positive lags mean that the fluctuations in the *iFV* signal precede those in that location. Negative lags mean the opposite, fluctuations appear in the voxel before they do in *iFV*. Lags vary vastly across tissue compartments (Figure 9.B; lags range [min/max=-11.04/9.28 s], [5^th^/95^th^ quantile=-9.15/-2.33 s]). Positive lags concentrate in ventricular regions superior to the *FV*, and negative lags are observed across grey matter. Right-left hemispheric symmetry is present, with longer negative lags appearing on most lateral regions (darker blue) as compared to more medial regions (light blue). Also, although significant correlations can be observed at many locations in the posterior half of the brain, the same is not true for frontal and inferior temporal regions. Next, figure 9.C shows cross-correlation traces in six representative voxels. We can observe large variability, with cross-correlation peaks occurring at different times in different locations.

**Figure 9.**
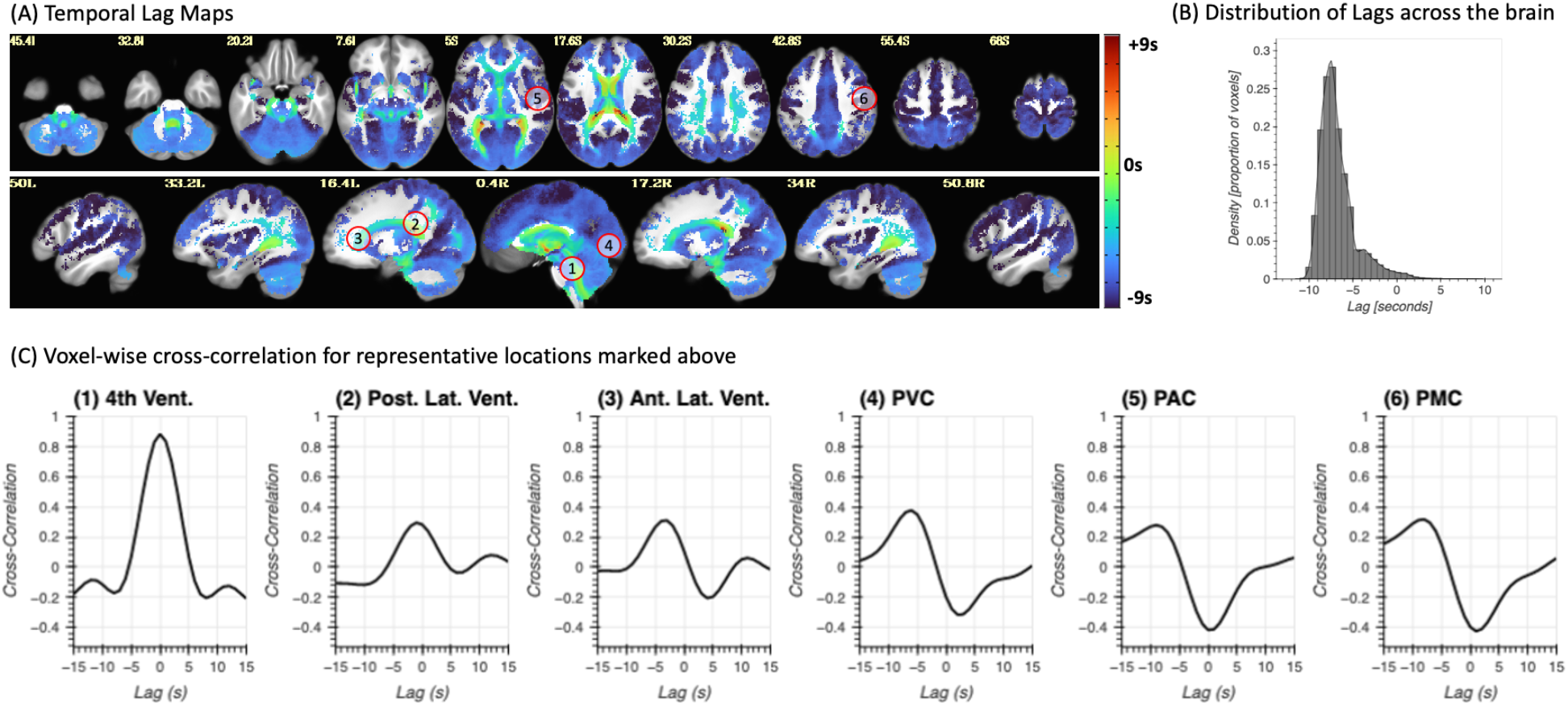
(A) Whole brain lag maps for the *iFV* signal. Color indicates the temporal lag conductive to maximal positive correlation within the lag range (−15s, 15s). A negative lag means the signal in that voxel precedes the *iFV*. (B) Distribution of temporal lags across the brain. (C) Representative voxel-wise cross-correlation traces for six different locations marked with numbered circles in panel (A).

### Effects on whole-brain functional connectivity matrices

Figure 10 shows the results for the whole-brain functional connectivity analysis. Both average connectivity matrices per scan type (e.g., “*awake*” or “*drowsy*”) and significant differences in connectivity between both types of scans were computed for three different pre-processing scenarios (i.e., *Basic, Compcor*, and *Compcor+*) to evaluate the effect that removal of ultra-slow fluctuations in *GM* time-locked with those in *iFV* may have on estimates of functional connectivity across the brain. Figure 10.A-E show results for the *Basic* pipeline where no signals from ventricular compartments are regressed. Figure 10.F-J show results when *CompCor* regressors (signals from all ventricles and WM) are included as additional regressors. Finally, Figure 10.K-O show results when voxel-wise optimally time-shifted versions of the *iFV* are also included as an additional regressor.

**Figure 10.**
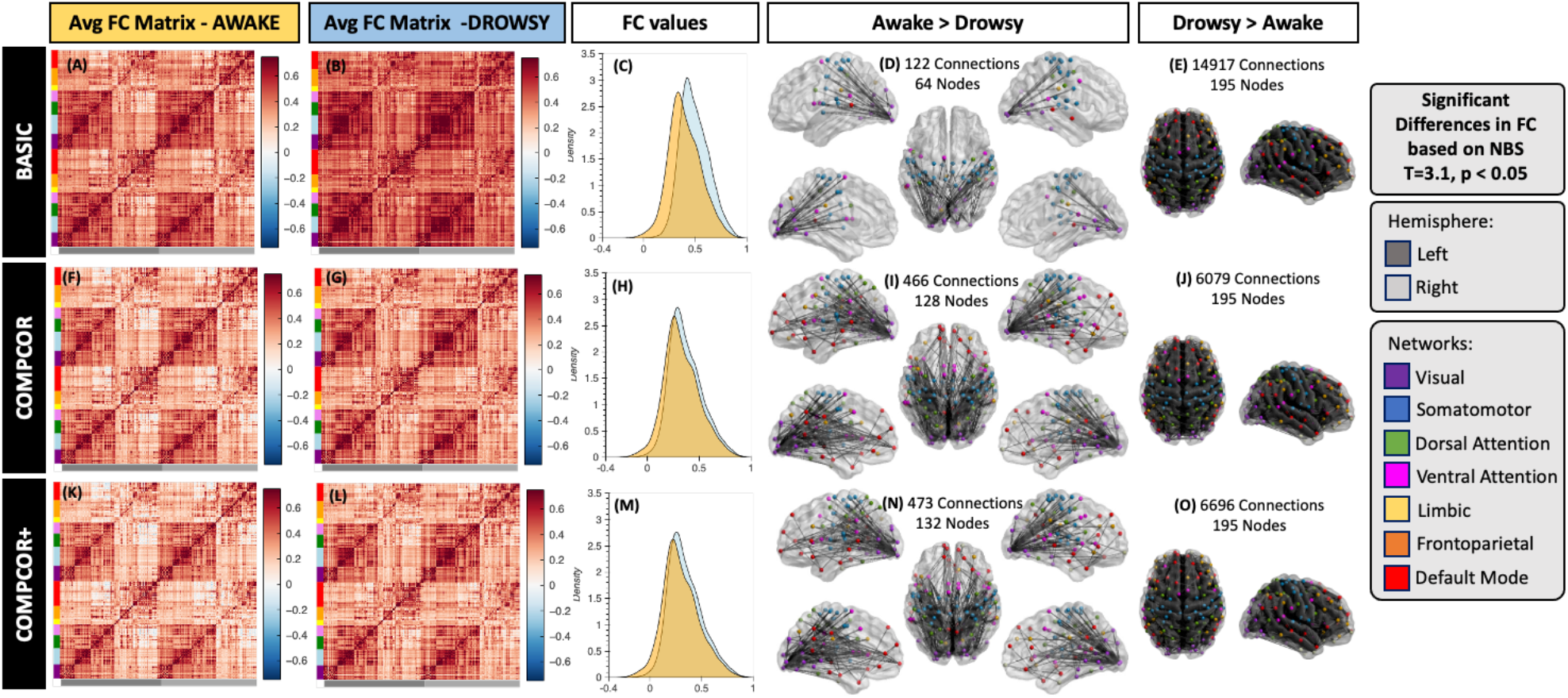
Whole-brain functional connectivity results. (A-E) Results for the *Basic* pre-processing pipeline. (F-J) Result for the *CompCor* pipeline. (K-O) Results for the *CompCor+* pipeline. In all rows the left most column shows the average connectivity matrix for all “*awake*” scans. The next column shows the average connectivity matrix for all “*drowsy*” scans. The middle column shows the distribution of correlation values across the whole brain for both scan types: drowsy in blue; and awake in orange. Finally, the last two columns show connections that were significantly different between the two scan types. The left most of these two columns show connections with stronger correlation for “*awake*” scans relative to “*drowsy*”. Conversely, the right most column shows those that are stronger for “*drowsy*” relative to “*awake*” scans. Significance was evaluated using Network-based statistics (T> 3.1, 5000 permutations, p<0.05 at the component level).

While average connectivity matrices for all three scenarios look quite similar for the “*awake*” scans (Figure 10.A, F & K); for “*drowsy*” scans we can observe stronger overall connectivity across most regions for the basic pipeline (Figure 10.B) compared to the two other pipelines that include as regressors signals from ventricular compartments (Figure 10.G & L). Additionally, no large differences in overall levels of connectivity values are observed between Figure 10.G (*CompCor*) and Figure 10.L (*CompCor+*) suggesting that the *CompCor* method accounts for a large portion of the ultra-slow fluctuations time-locked to the *iFV*.

The same trends are clearly observed in terms of the distribution of connectivity values across the whole brain presented in Figure 10.C, H & M. While for basic pre-processing (Figure 10.C) the distribution of connectivity values for “*drowsy*” scans is clearly shifted toward positive values relative to the distribution for “*awake*” scans. Such shift is not so apparent in the other two pre-processing scenarios (Figure 10.H & M).

When looking at significant differences in connectivity between “*awake*” and “*drowsy*” scans, we can observe the effects of using ventricular signals as nuisance regressors. First, the use of these regressors (both in the CompCor and CompCor+ pipelines) translates into higher sensitivity to detect connections that are stronger during “*awake*” scans compared to “*drowsy*” scans (Figure 10.D,I & N). Second, it also leads to large decrease in the number of connections rendered significant in the other direction (“*drowsy*” greater than “*awake”*; Figure 10. E, J & O). It is worth noticing that for the “*awake*” > “*drowsy*” comparison, significand differences in functional connectivity among frontal regions of the default mode network, and posterior parts of the brain only become apparent when ventricular signals are regressed during pre-processing.

For example, the percentage of connections with significantly stronger connectivity for “*drowsy*” compared to “*awake*” scans decreases from 79% (14917 connections) to 32% (6079 connections). Conversely, the percentage of connections that are significantly weaker for “*drowsy*” compared to “*awake*” subjects increases from 0.64% (122 connections involving 64 ROIs) to 2.5% (466 connections involving 128 nodes).

For comparative purposes, supplementary figure 3 shows results for the FC analyses when conducted on a variant of the Basic pipeline that also includes the *GS* as an additional nuisance regressors. In this case, average FC matrices for both conditions now contain a large proportion of negative correlations, concentrated primarily in connections between rather than within networks (panels a & b). Moreover, the distributions of connectivity values are centered around zero in this case (panel c). In terms of significant differences in FC across conditions, *GS* regression resulted also in higher sensitivity to detect connections that are stronger during “*awake*” scans compared to “*drowsy*” scans (3261 connections involving all ROIs), as well as a decrease in the number of connections rendered significant in the other direction (2971 connections involving all ROIs).

### How well can it predict periods of drowsiness?

#### Scan-level Results

To explore the potential value of *iFV* fluctuations around 0.05Hz as an indicator of lowered wakefulness, we decided to sort all 404 scans in terms of the total *PSD*_*sleep*_ (Figure 11.A). Because *GS*_*amplitude*_ has been previously stablished as a good indicator of wakefulness, we also sorted scans according to this second index (Figure 11.B). In figures 11.A & B, each scan is represented as a vertical bar whose height is either the *PSD*_*sleep*_ (10.A) or *GS*_*amplitude*_ (11.B). Each bar is colored according to whether that particular scan was labeled as “*awake*” (orange) or “*drowsy*” (light blue). In both plots, there is a higher concentration of “*drowsy*” scans on the left (i.e., higher *PSD*_*sleep*_ or *GS*_*amplitude*_). Conversely, there is a higher concentration of “*awake*” scans on the other end of the rank. This profile, of more “*awake*” than “*drowsy*” scans among those with higher *PSD*_*sleep*_ or *GS*_*amplitude*_, and the opposite as we go down on the rank becomes more apparent when we look at the relative proportions of each scan type in intervals of 100 scans (Figures 11.C & D). Within the top 100 ranked scans, the number of “*drowsy*” scans is more than double the number of “*awake*” scans. Conversely, we look at the bottom 104 ranked scans, these proportions have reversed. Finally, it is worth noticing that *PSD*_*sleep*_ and *GS*_*amplitude*_ based rankings, although similar at the aggregate levels described here, are not equal on a scan-by-scan basis (see suppl. figure 9 for a scatter plot of scan-level *PSD*_*sleep*_ vs. *GS*_*amplitude*_). In other words, the specific rank of a given scan is not necessarily the same for both ranking schemes.

**Figure 11.**
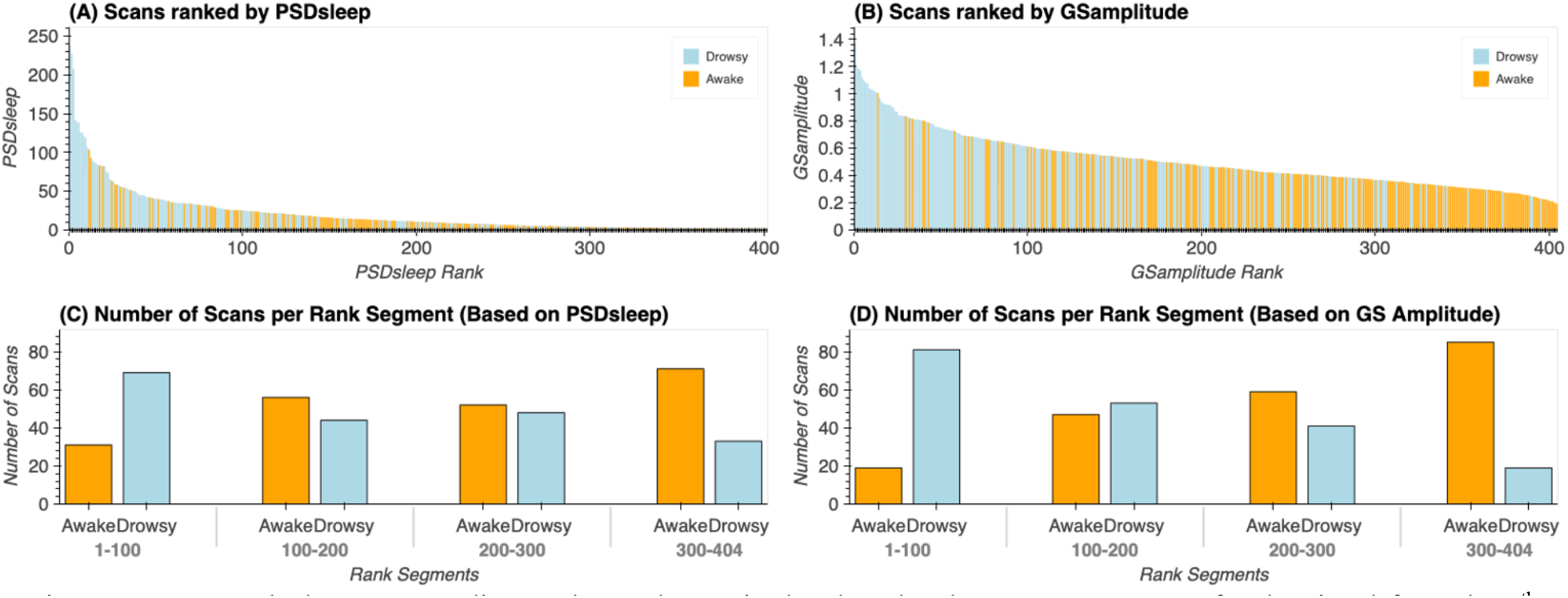
(A) Ranked scans according to the total PSD in the sleep band (0.03Hz – 0.07Hz) for the signal from the 4^th^ ventricle. (B) Ranked scans according to the amplitude of the global signal. In both (A) and (B), each scan is represented by a vertical bar. The color of the bar indicates whether a given scan was labeled “drowsy” or “awake” based on the total amount of time subjects kept their eyes closed during that particular scan. (C) Proportion of “drowsy” and “awake” scans in rank intervals (1 – 100, 101 – 200, 201 – 300, 301 – 304) for PSD-based ranks. (D) Proportion of “drowsy” and “awake” scans in rank intervals (1 – 100, 101 – 200, 201 – 300, 301 – 304) for GS-based ranks.

#### Segment-Level Results

We also evaluated the ability of *PSD*_*sleep*_ to predict wakefulness at the segment level (i.e., 60 seconds segments), and how it compares to that of *GS*_*amplitude*_. Figure 12 shows results for this last set of analyses that look at prediction accuracy under several scenarios: a) when *PSD*_*sleep*_ is the only input feature, b) when *GS*_*amplitude*_ is the only input feature (we attempt this for all pre-processing pipelines separately), c) when both *PSD*_*sleep*_ and *GS*_*amplitude*_ are used as input features (*PSD+GS*) and d) when *PSD*_*sleep*_ is the only feature and labels are randomized (control condition). Accuracy was significantly higher (T-test; p<1e^-6^ in all instances) for the three non-control scenarios (i.e., *PSD*_*sleep*_ : 71%, *GS*_*amplitude*_ : 74%, *PSD+GS: 70%)* when compared to the control condition (Control: 40%). Although significant differences were also observed among the three non-control scenarios (not marked in figure 12.A), in this case, differences in accuracy in absolute terms was very small (<0.04). No significant difference in accuracy exists for *GS*_*amplitude*_ computed following the different pre-processing pipelines.

**Figure 12.**
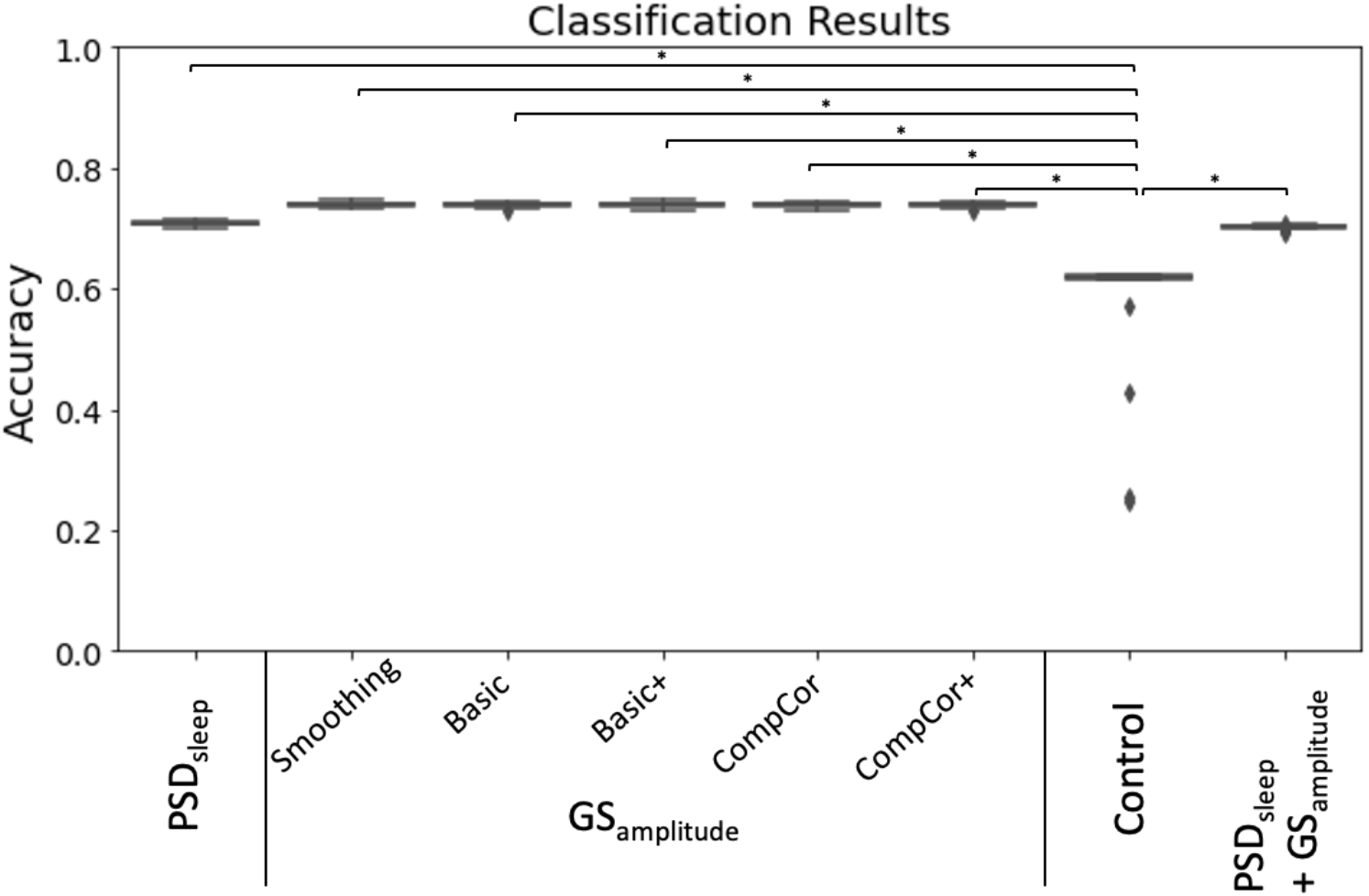
Prediction accuracy for a logistic regression classifier with different set of input features: PSD_sleep_, GS_amplitude_ (computed for different pre-processing pipelines), randomized PSD_sleep_ (control) and PSD_sleep_ + GS_amplitude_.

## Discussion

Previous work has shown that strong ultra-slow (∼0.05Hz) inflow fluctuations appear in the *FV* when subjects descend into sleep, and that anticorrelated BOLD fluctuations of equal frequency can be observed in grey matter (Fultz et al., 2019). A similar observation was made earlier this year for awake scans (Yang et al., 2022). Here, we reproduce those findings in a much larger sample (174 subjects here vs. 13 in (Fultz et al., 2019) and 10 in (Yang et al., 2022)), and then we extend them in several meaningful ways.

### Nature of iFV ultra-slow fluctuations and relationship to the GS

Ultra-slow *FV* fluctuations (∼0.05Hz) with inflow properties (i.e., decreased intensity across successively acquired slices) were detected in the 7T HCP sample, despite this data not having optimal sensitivity for inflow effects at this location. Additionally, fluctuations in *iFV* showed maximal negative correlation to the *GS* for a positive lag of 2s (same as in Fultz et al.), and maximal positive correlation with *-dGS/dt* for a negative lag of −3s (−1.8s for Fultz et al.). Agreement between the observations made here and those in Fultz et al. suggest we are looking at signals of similar nature, namely CSF inflow into the *FV* driven by a decreases of cerebral blood volume (as indexed by the *GS*) and the need to keep the sum of all brain volumes inside the skull constant (Monro-Kellie doctrine; (Mokri, 2001)).

If so, then how can we explain the presence of residual inflow effects on a dataset where the *FV* sits far away from the lower end of the imaging FOV and TR is not as sort (1s here vs. 367ms in Fultz et al.)? First, sensitivity for inflow increases with field strength and flip angle (Gao and Liu, 2012); both of which are higher here compared to Fultz et al. (2019). Second, in the HCP data, the *iFV* sits, on average, somewhere between slice 17 and slice 24. Given the use of a multi-band sequence (Feinberg and Setsompop, 2013) with factor (MB=5) and a number of slices equal to 85 in the 7T HCP protocol, the minimal critical imaging velocity (Fultz et al., 2019; Kim and Parker, 2011) for some slices sitting over the *FV* is 54.40mm/s. Awake CSF maximal velocity ranges between 20-80mm/s in healthy adults (Korbecki et al., 2019; Lee et al., 2004; Nitz et al., 1992). Although the velocity for reversed CSF flow during sleep might be different, Fultz et al. already reported fast events observable four slices away from the lower boundary of the FOV, which would require, for their protocol, CSF velocities above 68mm/s. These numbers show that inflow effects may be still observable in the HCP data, even if the location of the slices and the TR is not optimally sensitized. The same might be true for other data acquired with multi-band protocols, and additional research should elucidate how often these fluctuations are present in resting-state datasets. This is important, given one can expect a third of subjects to fall asleep within 3 minutes of scan onset (Tagliazucchi and Laufs, 2014), and that—as we shall discuss below—these fluctuations can significantly affect estimates of functional connectivity.

Examination of slice-dependent intensity profiles revealed two different patterns within the *FV* (figure 2). Slices inferior to the *pDmR* exhibited an inflow-like profile with decreased signal intensity across successively acquired slices. Conversely, for slices superior to the *pDmR*, CSF signal intensity decreases moving rostrally towards the brain. Despite its small size, the *FV* is a complex structure with openings towards several CSF compartments, namely the third ventricle via the aqueduct of Sylvius, the central canal via its narrowing at the Obex, the cisterna magna via the median aperture (or foramen of Magendie) and the pontocerebellar cistern via the two lateral apertures (of foramina of Luschka). As such, its fluid dynamics are complex. In our sample, it seems that inflow effects due to fresh CSF coming from outside the imaging FOV are only dominant for signals recorded on the caudal half of the *FV* before such fresh fluid mixes with that coming from all other chambers that sit inside the imaging FOV. Based on this observation we decided to constrain our analyses to *FV* signal extracted using only the caudal half of the *FV* (what we refer to as the inferior *FV* or *iFV* throughout the manuscript).

Here we decided to look beyond global effects (e.g., relationships to the *GS*) and explore in detail the spatio-temporal profile of ultra-slow BOLD fluctuations time-locked to those in the *iFV*. While doing so, we observed that the strongest anticorrelations between those two signals in drowsy scans occurred in visual, *MT*, sensorimotor, and supplementary motor cortexes (figure 8). This pattern overlaps with that previously reported by Song et al. (2019) for fluctuations of similar frequency that appear in fMRI recordings during light sleep and track EEG spindles (a key signature of light sleep; (Gennaro and Ferrara, 2003)). Song et al. (2019) also reported that these oscillations first appear in the thalamus and sensory regions; and from there, they propagate to prefrontal regions. Here, we partially observe a similar pattern with sensory regions (e.g., motor, auditory cortex) having larger negative lags (e.g., leading more in front of the *iFV*) than frontal regions (figure 9.A, supp. figure 7). These converging observations emerge from a very different set of methods and assumptions. In the case of Song et al., the authors relied on concurrent EEG measures to detect periods of light sleep and subsequently evaluate changes in frequency profiles at the regional level relative to wakefulness. Here, we simply studied patterns of correlation with the *FV* signal on scans marked as “*drowsy*” based on eye tracking recordings. Once more, agreement across studies suggest that the bulk of our observations are indeed related to periods of light sleep, and not simply to subject’s lack of compliance regarding instructions to keep their eyes open. This is important, as with ET alone it is not possible to separate these two potential scenarios.

### Contribution to GSamplitude during drowsiness

It is well established that the standard deviation of the global signal (*GS*_*amplitude*_) increases when vigilance decreases (Wong et al., 2013) and when subjects fall asleep (Fukunaga et al., 2006; Horovitz et al., 2008; Larson-Prior et al., 2009). In agreement with these prior findings, here we observed a significantly higher *GS*_*amplitude*_ for long segments of EC compared to EO (Figure 5.A). Importantly, these differences remain significant for all other pre-processing pipelines, including those that attempt to remove *iFV* related fluctuations (supp. table 1). This suggests that the ultra-slow sleep-related fluctuations being studied here are not the sole driver of differences in *GS*_*amplitude*_ across wakefulness states. This may be initially counterintuitive given those sleep-related fluctuations are quite strong and widely distributed in space. Yet, for any spatially distributed signal to be a contributor to the *GS*, such signal ought to appear in synchrony across the brain. Otherwise, positive and negative contributions from different locations may cancel each other (see (Liu et al., 2017) for a detailed description of the *GS* as a measure of synchronicity across the brain). Our results suggest that the reported heterogeneity regarding temporal lags (Figure 9) results in an overall effect that, although important, does not fully explain the increase in *GS*_*amplitude*_ that accompanies decreases in wakefulness. This is not to say that such fluctuations have no effect at all. As supplementary table 1 shows, differences in *GS*_*amplitude*_ between EC and EO segments decrease substantially for pre-processing pipelines that try to account for signals that correlate with those present in the ventricles (including the *FV*), yet not to a point of eliminating significance. Ultimately, what this suggests, is that differences in *GS*_*amplitude*_ should not be attributed only to physiological effects that create an imprint in ventricular regions (e.g., cardiac, respiration, sleep-related inflow), and that other effects constrained to grey matter are also important.

### Ability to predict drowsiness

The primary goal of this work was to evaluate how well spectral characteristics of ultra-slow inflow fluctuations in *iFV* could help us predict drowsiness during resting-state scans at three different levels: sample level (i.e., the whole dataset), scan level (i.e., that of individual scans), scan-segment level (i.e., that of portions of individual scans).

Regarding sample-level, it has been previously shown that approximately a third of subjects in the 3T resting-state HCP sample fell asleep within three minutes from scan onset (Tagliazucchi and Laufs, 2014). Here we report on a similar pattern for the 7T HCP sample (Figure 7). While that earlier report on the 3T sample relied on a hierarchical tree of pre-trained support linear vector machines, here we base our observation on the progressive increase in *PSD*_*sleep*_ for the *iFV* signal. For the initial 180 seconds of scanning, average *PSD*_*sleep*_ traces for “*awake*” and “*drowsy*” scans overlap. Beyond that temporal landmark, *PSD*_*sleep*_ for “*drowsy*” scans rises above that of “*awake*” scans. Conversely, the *PSD* for the control band (which excludes the 0.05Hz signals of interest) did not differ across scan types irrespective of time; demonstrating the specificity of the effect to the “*sleep*” band. It is worth noticing that these patterns are clearly observed despite the large confidence intervals associated with across-scans averaged *PSDs* (Figure 5.A); which emanate from the fact that subjects fall asleep at different moments. Yet, because we can expect a monotonic increase in the number of subjects that fall asleep as scanning time increases, a clearer pattern emerges when looking at cumulative PSDs (Figure 5.B). While the cumulative *PSD*_*control*_ for “*drowsy*” and “*awake*” scans have equivalent small increases with time; *PSD*_*sleep*_ traces clearly diverge across scan types, with “*drowsy*” scans showing a much faster increase, particularly after approx. 200 seconds from scan onset. These results demonstrate that an easily implementable metric, such as the *PSD*_*sleep*_ of the *FV*, can provide equivalent information about the progressive descent into sleep of a resting-state sample to that obtained using relatively more complex machine learning procedures.

Next, our scan-level exploration showed that sorting scans based on average *PSD*_*sleep*_ results in a list (Figure 11.B) with “*drowsy*” scans appearing in higher proportion (69 drowsy / 31 awake) at the top of the rank (e.g., higher average *PSD*_*sleep*_ levels), and “*awake*” scans appearing at higher proportion (71 awake / 33 drowsy) at the bottom of the rank (e.g., lower average *PSD*_*sleep*_ levels). Unfortunately, the rank is not perfect, meaning there is no certainty that a particular scan near one end of the ranking (e.g., scan ranked 400/404) is of one specific type (e.g., “*awake*”). Several reasons contribute to this situation. First, the dichotomic labeling system (“*drowsy*” / “*awake*”) based on the presence/absence of long periods of eye closure may be too simplistic to represent the heterogeneity of sleep patterns present in the data. Similarly, a single number (average *PSD*_*sleep*_) may not suffice to characterize the wakefulness profile of complete scans with alternating periods of wakefulness and sleep of different durations and intensity. Second, while the absence of long periods of eye closure can be confidently associated with wakefulness, the presence of long periods of eye closure does not necessarily mean sleep. As such, some scans labelled here as “*drowsy*”, may not include sleep periods, and simply capture the fact that subjects stopped complying with the experimental request to keep their eyes open. Finally, another contributing factor might be that not every instance of increase *PSD*_*sleep*_ is necessarily associated with a period of light sleep. A closer look at some of the ranking “*misplacements*” at the top of the rank—namely “*awake*” scans with high average *PSD*_*sleep*_ (supp. figure 8) shows that in some instances (subjects A and B) periods of elevated *PSD*_*sleep*_ coincide with peaks of head motion (black arrows); yet that is not always the case (blue arrows). Although we did not find a significant difference in head motion across scan types, examples such as these suggest that strong ultra-slow fluctuations in the *iFV* can indeed sometimes appear due to motion, decreasing the accuracy of *PSD*_*sleep*_ as a frame-to-frame marker of sleep at the individual level.

We also observe periods of elevated *PSD*_*sleep*_ that do not overlap with periods of higher motion or eye closure (e.g., subjects C & D in supp. figure 8). This suggests that fluctuations at around 0.05Hz in the *FV* are not exclusive to light sleep and can also occur during wakefulness in the absence of motion. This agrees with observations by Yang et al. (2022), who just reported the presence of ultra-slow inflow fluctuations in the *FV* in a sample of awake resting-state scans. Yang and colleagues suggest that although similar in their presentation, ultra-slow *FV* inflow fluctuations are driven by cerebral blood volume oscillations of different origin: neuronal during sleep and due to changes in blood gases concentrations (i.e., pCO_2_) during awake states. As both phenomena can influence *PSD*_*sleep*_ similarly, this is a likely important contributor to ranking errors and segment misclassification (discussed in the next paragraph).

Finally, our segment-level results show that *PSD*_*sleep*_ computed on 60 seconds segments can predict wakefulness (e.g., eyes open/awake vs. eyes closed/drowsy) with 71% accuracy (figure 12), slightly less than what is attainable (74%) using *GS*_*amplitude*_ as the input feature for classification. The value of *GS*_*amplitude*_ as a marker of vigilance/arousal is well stablished. Our result suggests that the spectral content around 0.05Hz of the *FV* signal (i.e., *PSD*_*sleep*_) can be equally informative. When the classifier is given both metrics as input features (*GS*_*amplitude*_ and *PSD*_*sleep*_), accuracy levels barely changed suggesting these two metrics did not provide complementary information the logistic classifier could exploit to increase accuracy. As originally stated, our goal was to keep methods as simple as possible, and that was the reason why we chose a simple logistic regressor for this set of analyses. It is possible that more advanced classification algorithms (e.g., support vector machines, neuronal networks) might be able to find complementary bits of information in both traces, and therefore the fact that the accuracy for a logistic regression machine barely improved when both metrics are combined should not be taken as evidence that the same will occur with other models.

### Effects on functional connectivity

Regression of ventricular fluctuations (as in *CompCor* and *CompCor*+) had an equalizing effect on the distribution of FC estimates across the brain for “*drowsy*” and “*awake*” scans (figure 10.H & M). It also resulted into increased sensitivity and specificity for detecting FC differences between those two wakefulness states (Figure 14.D, E, I and J) and better agreement with previous findings regarding connectivity changes during eye closure and sleep.

When ventricular signals are not included as nuisance regressors, comparison of connectivity patterns for “*drowsy*” and “*awake*” scans produced two main observations: a) the number of connections with increased connectivity during “*drowsy*” scans is much larger than those that show a decrease in connectivity; and b) most decreases in connectivity for “*drowsy*” scans involve visual regions. These observations persist if ventricular signals are used as nuisance regressors during pre-processing. Yet, additional observations become possible such as decreased intra-network connectivity for “*drowsy*” scans in the visual, attention and default mode networks, as well as between nodes of the visual network and frontal eye field regions from the dorsal attention network. All these observations agree with those recently reported by Agcaoglu et al. (2019) when comparing eyes open to eyes closed resting-state scans. One key discrepancy between our results that those of Agcaoglu et al., is that we also observed decreased connectivity between motor and visual regions in “*drowsy*” scans. Yet, such trend has also been previously reported by Van Dijk and colleagues (2010) when comparing eyes open vs. eyes closed; suggesting this pattern is not unique to this dataset. Among the changes only observable when ventricular signals are modeled as nuisance regressors, we encounter a decrease in connectivity between anterior and posterior nodes of the default mode network, which is a key signature of sleep (Horovitz et al., 2009). Similarly, we also observe decreased connectivity for regions of the dorsal and ventral attention network. Reduced connectivity in fronto-parietal networks during sleep is another common finding in the literature (Larson-Prior et al., 2009; Picchioni et al., 2013) that is consistent with a behavior, sleep, that is characterized by reduced attention towards the external environment. Overall, these results suggest that removal of ultra-slow fluctuations is key to properly identifying previously described changes in functional connectivity that accompany eye closure and sleep. Yet, this does not necessarily imply that such an approach is free of interpretational confounds. If, as Yang and colleagues suggest, ultra-slow BOLD fluctuations time-locked to those in *iFV* have different etiology during awake and drowsy states, modeling these fluctuations as nuisance regressors may inadvertedly remove neural activity that varies between awake and drowsy states. Moreover, given both our results, and prior reports (Tagliazucchi and Laufs, 2014) demonstrate that one can expect a range of drowsiness states in any resting-state sample, the same regressor may be modeling different effects (i.e., neuronal or physiological) in different parts of the sample. Given those considerations, researchers should always consider the acquisition of concurrent physiological recordings (e.g., pulse oximeter, respiration belt (Birn et al., 2006; Glover et al., 2000), peripheral optical imaging (Tong et al., 2015, 2013)) in fMRI samples likely to include fluctuations in wakefulness (e.g., resting-state).

When the *GS* is used as an additional regressor (supplementary figure 3.A-G) we observe a shift in the distribution of correlation values towards zero, which consequently resulted in the appearance of negative correlations that were not present otherwise. Prior work has demonstrated that *GS* regression (GSR) mathematically imposes such a shift and the appearance of negative correlations (Murphy et al., 2009; Murphy and Fox, 2017). In terms of differences across states (“*drowsy*” vs. “*awake*”), GSR produced substantially different results to all other pipelines. All non-GSR pipelines are characterized by a much larger number of connections being significantly stronger for “*drowsy*” states. For the “*awake > drowsy*” contrast, connections primarily included ROIs in the visual network as well as previously discussed signatures of sleep in DMN and attention networks. Following GSR, the number of significantly different connections in both directions was more similar, and the “*awake > drowsy*” connections were now distributed across most networks, in addition to the connections observed without GSR (supplementary figure 3.D-F & H). This is likely a result of documented observations that GSR will distribute distinctive signal fluctuations in focal brain regions into other regions (Aguirre et al., 1998) and will distribute connections that are focal to a subset of ROIs across other brain regions (Gotts et al., 2020, 2013; Saad et al., 2012). Also, these results highlight that the regression of a signal component (i.e., the *GS*) with strong links to vigilance (Liu et al., 2017) can alter observed differences in connectivity across scans with differing levels of drowsiness.

### Study limitations

First, we cannot completely rule out the confounding effects of differences in cardiac and respiratory function across wakefulness states. Cardiac traces estimated directly from the fMRI data showed there is a small, but significant decrease in cardiac rate for periods of elevated drowsiness. Although such differences dissipated when looking at their aliased equivalent based on the fMRI sampling rate, we cannot rule out that second order effects (e.g., cardiac rate variability (Chang et al., 2013)) may be a contributing factor to observed differences. In addition, as no respiratory traces are available, we cannot evaluate the role of respiration to these results. Second, as previously discussed, we indirectly infer periods of sleep/drowsiness based on eye closure. Although we took actions to remove from the analysis long periods of eye closure that were likely the result of subject’s lack of compliance with instructions to keep their eye open, we acknowledge that our proxy for sleep/drowsiness is not as accurate as an EEG based measure. Future work can use a large sample size of multi-modal EEG/fMRI data to better validate our current observations.

Third, we infer an inflow origin for the targeted *iFV* fluctuations based on their across-slice profile and their relationship to cerebral blood volume as indexed by the GS and its first derivative. Yet, alternative ways to test the inflow nature of this phenomena should be explored in future studies. For example, one could rely on multi-echo fMRI—which refers to the concurrent acquisition of fMRI at multiple echo-times (Posse et al., 1999)—to separate signal fluctuations of BOLD (not inflow-related) and non-BOLD (inflow-related) origin based on how their amplitudes are modulated by the different echo times (Caballero-Gaudes et al., 2019; Gonzalez-Castillo et al., 2016; Kundu et al., 2011). One could also acquire fMRI data at multiple flip angles or repetition times (TR). These two imaging parameters affect the amount of T1-weigthing present on the recorded signals, and as such would have a strong modulatory effect on inflow-related fluctuations (Gao and Gore, 1994; Gao and Liu, 2012). Similarly, one could acquire data using two different slice geometries: one with slices parallel to the hypothesized direction of flow (e.g., sagittal slices), and one perpendicular to it (e.g., axial slices). Inflow-related fluctuations for the hypothesized direction of flow should only be present in the perpendicular acquisition (Gao and Liu, 2012). Finally, researchers could also use outer volume saturation techniques to attempt nulling fluctuations due to incoming fluid from anatomical regions right underneath the imaging field of view.

## Conclusions

Resting-state data is a chief component of today’s non-invasive human neuroimaging research (Essen et al., 2013; Miller et al., 2016). Despite its long history (Biswal, 2012; Snyder and Raichle, 2012), there are many open questions regarding what mechanisms lead to the formation of resting-state networks and their time-varying profiles (Gonzalez-Castillo et al., 2021). One such question is the role that shifts in vigilance and arousal play in shaping resting-state results (Laumann et al., 2016; Tagliazucchi and Laufs, 2014), which remains rarely explored because it requires concurrent electrophysiological or eye tracking recordings. Here we study how signatures of CSF inflow during light sleep in fMRI data (Fultz et al., 2019) can be exploited to extract information about wakefulness levels in resting-state data. Our results demonstrate that it is possible to detect this phenomenon in data with relatively small inflow weighting, and that it has value as a marker of wakefulness fluctuations at the sample, scan and segment levels. We also discuss how the use of ventricular signals as nuisance regressors significantly alters inferences about how FC changes during sleep because of the link (i.e., co-linearity) between ultra-slow fluctuation of inflow origin in the FV and those of BOLD origin in GM. These considerations are important because, as we reproduce here for the HCP 7T dataset, large resting-state samples can often include scans from subjects with quite different wakefulness profiles. Today, in most resting-state studies, those differences are ignored, and all resting-state scans are considered equivalent. This can translate into the undesired mixture of two distinct connectivity profiles into one that may not be easily interpretable, as it does not represent the average of a homogenous sample (see (Liu and Falahpour, 2020) for a discussion of a similar concern specific to vigilance). Data-driven methods such as the one described here, allow us to easily extract information about wakefulness states, and should help ameliorate this issue by helping us partially segregate scans and segments into more homogenous samples.

## Acknowledgements

This research was possible thanks to the support of the National Institute of Mental Health Intramural Research Program (ZIAMH002783). Portions of this study used the high-performance computational capabilities of the Biowulf Linux cluster at the National Institutes of Health, Bethesda, MD (*biowulf*.*nih*.*gov*).

