## Supplementary Materials for "Ultra-slow fMRI fluctuations in the fourth ventricle as a marker of drowsiness"

### Supplementary Figures

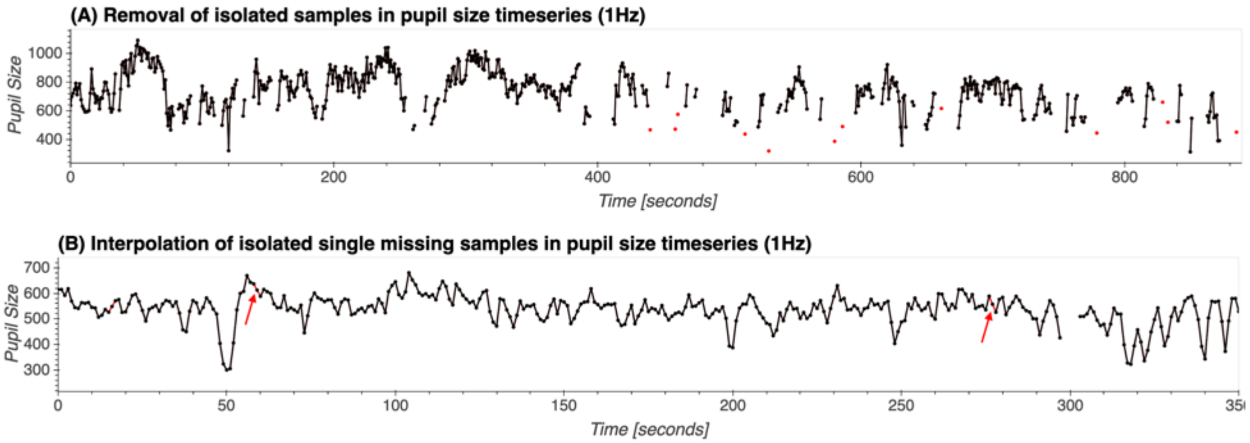


Supplementary Figure 1. Representative cases of final corrections to eye pupil timeseries once those have been interpolated to a sampling rate of 1Hz. (A) In some cases, following interpolation, we observed the presence of isolated samples scattered during long periods of eye closure (red dots). If those samples are not removed, these long periods of eye closure would not be identified as such. To avoid that, we removed any isolated sample with missing data on both ends. (B) Similarly, we observed individual gaps of a single sample breaking long periods of eye closure (red arrow). In those instances, those missing samples were interpolated (red trace).

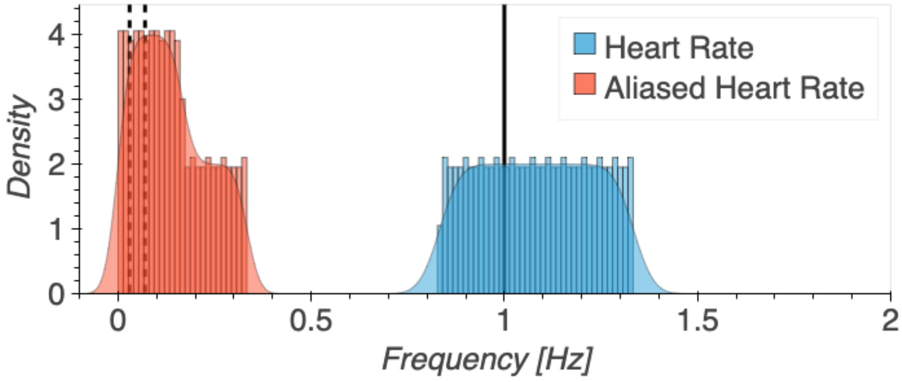


Supplementary Figure 2. Simulation of frequency aliasing for cardiac pulsations. The sampling frequency of the fMRI data is 1Hz (black continuous line). Our target fluctuations of interest sits around 0.05Hz, and to detect it we focus our attention on the frequency range [0.03Hz - 0.05 Hz] (narrow band between the two vertical black dashed lines). Typical cardiac rates range from 50 to 80 beats per minute while subjects are resting (blue histogram/distribution). Due to frequency aliasing, cardiac pulsations at those frequencies will appear at lower parts of the spectrum in the fMRI recordings. As the figure shows, for a sampling frequency Fs=1Hz there is potential for those to overlap (red histogram/distribution) with the target frequency of our study.

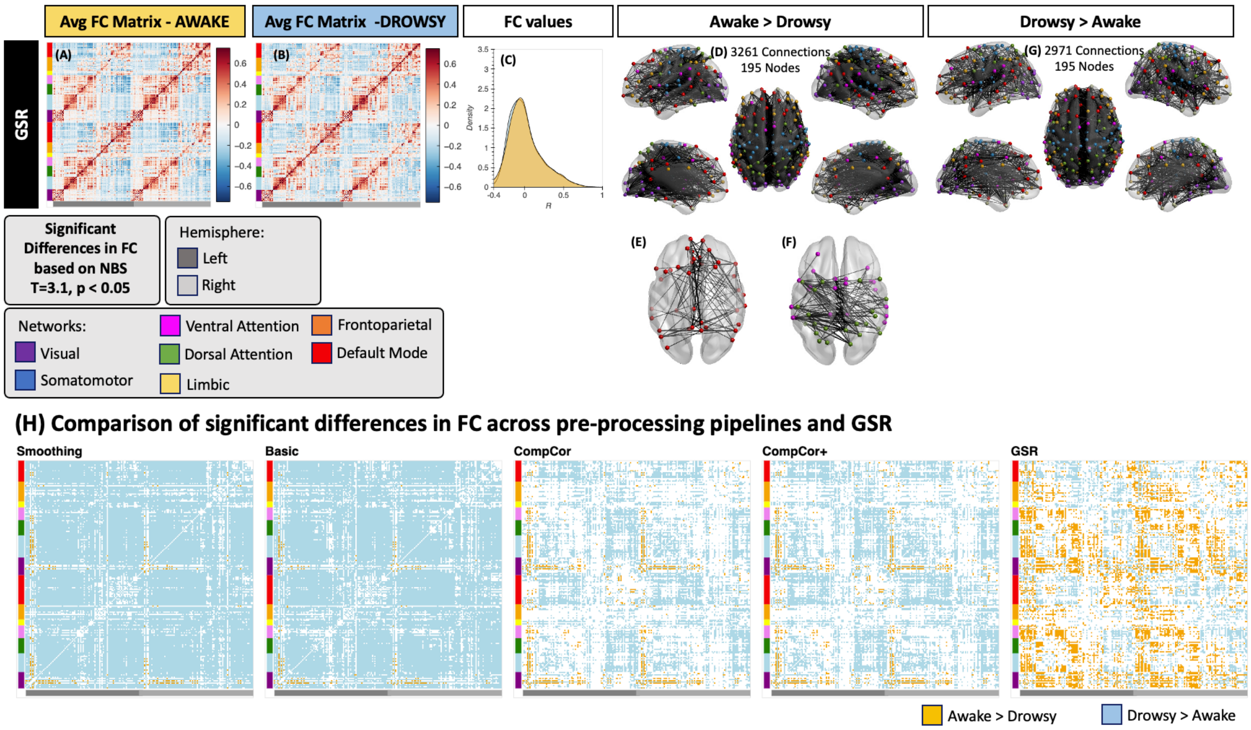


Supplementary Figure 3. Significant differences in FC when GS is used as an additional nuisance regressor in the Basic Pipeline. (A) Average connectivity matrix for all "*awake*" scans. (B) Average connectivity matric for all "*drowsy*" scans. (C) distribution of correlation values across the whole-brain for both scan types: drowsy in blue, awake in orange. (D) All connections that appear as significantly stronger for "*awake*" compared to "*drowsy*" scans when performing global signal regression (GSR). (E) Within default-mode network connections that appear as stronger for "*awake*" compared to "*drowsy*" scans. These are a subset of the connections shown in (D). (F) Similar to (E), but this time we show connections within both attentional networks. (G) All connections that appear as significantly stronger for "*drowsy*" compared to "awake" scans when performing GSR. (H) Significantly different connections in both contrast directions ("*awake > drowsy*" in orange, "*drowsy > awake*" in blue) for all considered pre-processing pipelines.

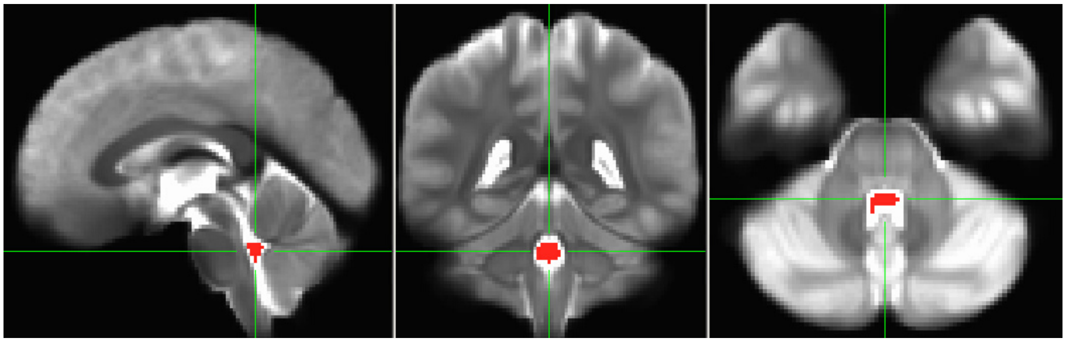

Supplementary Figure 4. Region of interest for the inferior 4th ventricle overlaid on top of the average of the first EPI image from all 404 resting state scans entering the main analyses.

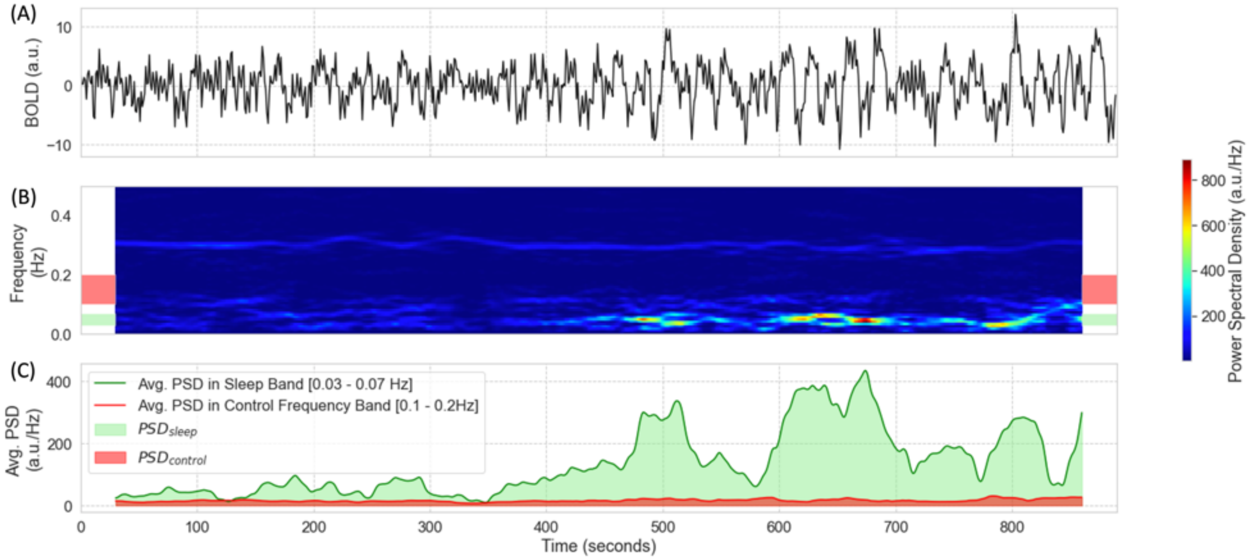


Supplementary Figure 5. Evaluation of the temporal evolution of fluctuations around 0.05Hz in the BOLD signal from the *iFV* in one representative subject. (A) BOLD timeseries for the 4th ventricle during one representative 15 minutes resting state scan. (B) Spectrogram for the representative time-series depicted in the top panel. Two frequencies bands of interest are marked as colored bands on the sides of the spectrogram: 0.03 – 0.07Hz (sleep band – green shading) and 0.1 – 0.2 Hz (control band – red shading) (C) Average PSD for the two frequency bands of interest: sleep band (green) and control band (red). Bold lines represent the temporal evolution of the average PSD for each band. Shaded area, namely the area under a given PSD trace, represents the total PSD in each band for the whole scan.

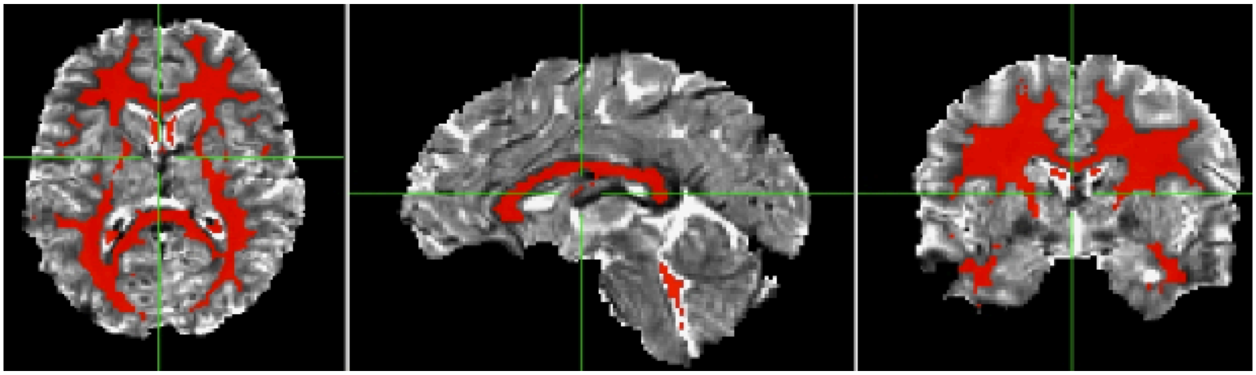

Supplementary Figure 6. Region of interest for the *CompCor* method overlaid on top of the average timeseries for one representative fMRI scan.

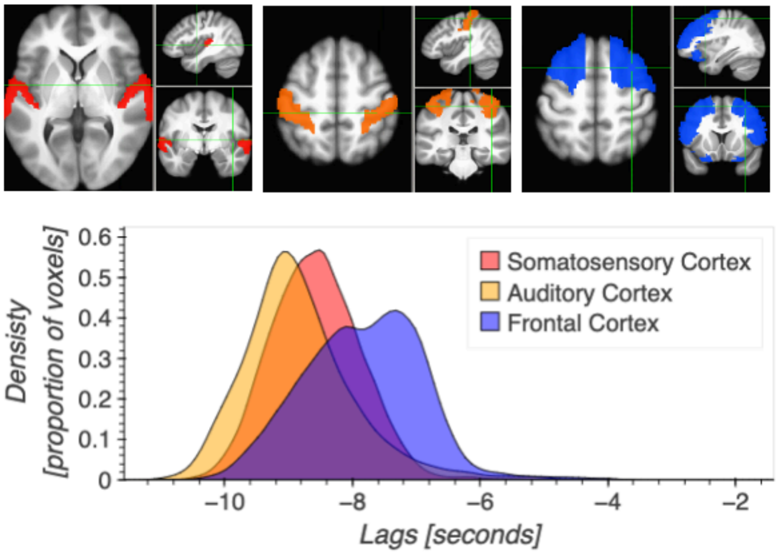


Supplementary Figure 7. Distribution of lags for three macro-anatomical regions: somatosensory cortex (red), auditory cortex (orange), frontal cortex (blue). The top row shows the three ROIs, which were defined using the version of the Eickhoff-Zilles atlas distributed with the AFNI software. The bottom row shows the distribution of temporal lags for these three regions. One can observe that fluctuations time-locked to the *iFV* appear in somatosensory and auditory cortex prior to frontal cortex.

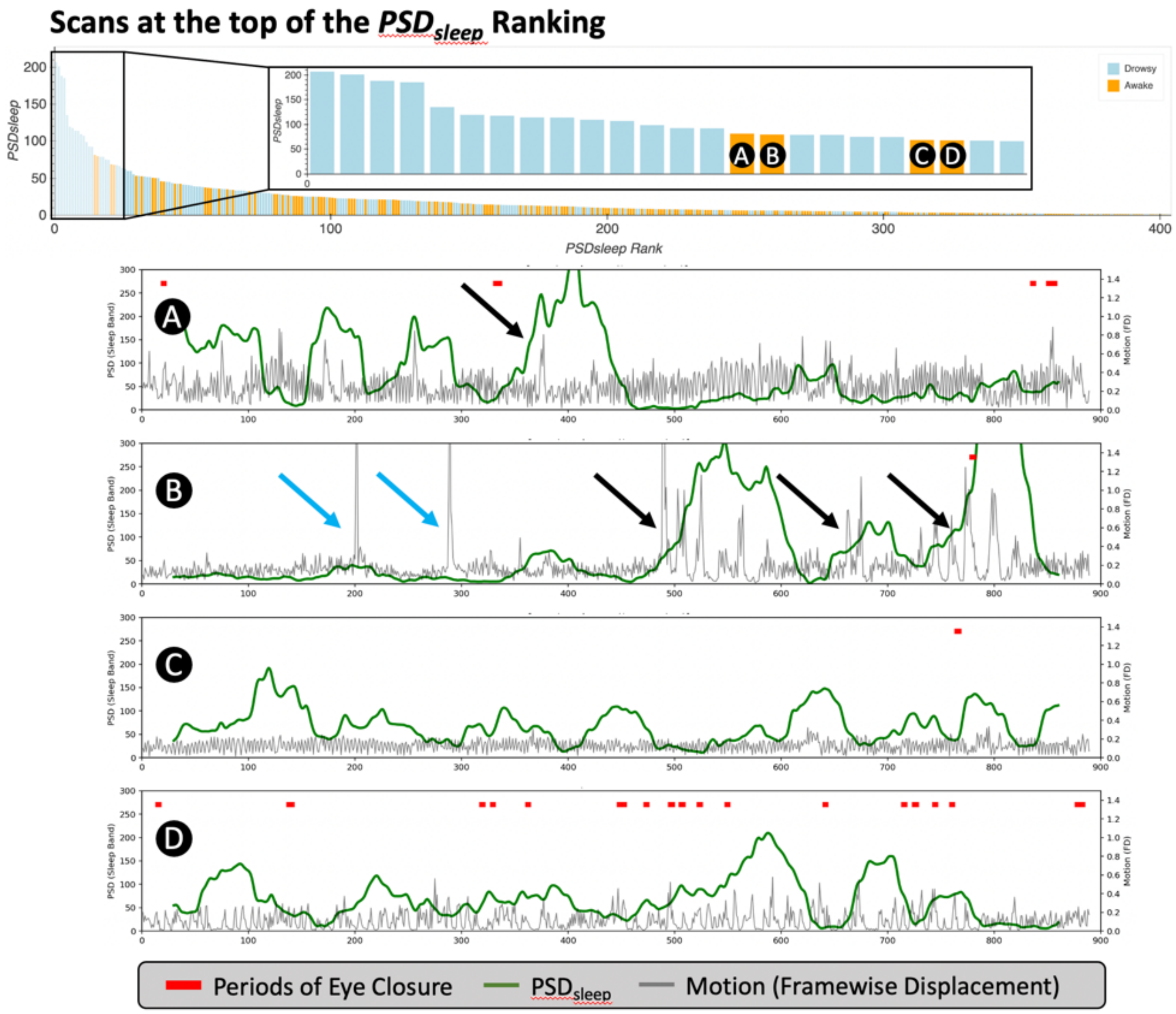


Supplementary Figure 8. The top panel shows a zoom-in version of the scan rank based on $PSD_{sleep}$ (Figure 11.A). The first four "miss-ranked" subjects are marked as A, B, C and D. Those are "awake" scans that show a high level of $PSD_{sleep}$. Below this first panel, we show traces of $PSD_{sleep}$ (green) and framewise displacement (grey) for the four subjects. Additionally, we mark periods of eye closure with red blocks. Black arrows indicate instances where increases in $PSD_{sleep}$ are accompanied by spikes in head motion. Blue arrows show instances of head motion that are not accompanied by an increase in $PSD_{sleep}$. Finally, subjects C & D are good examples of subjects that show periods of elevated $PSD_{sleep}$ in absence of eye closure or motion spikes.

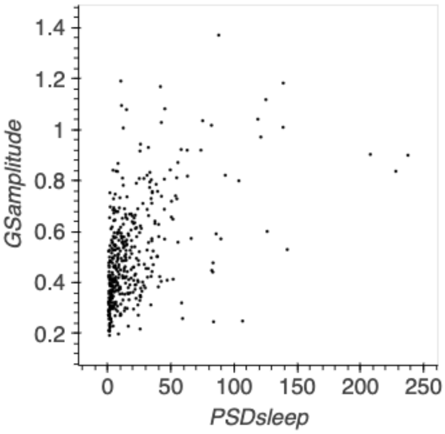


Supplementary Figure 9. Scatter plot of scan-level average $PSD_{sleep}$ versus the amplitude of the global signal.

### Supplementary Tables

| Preprocessing Pipeline | Eyes Closed | Eyes Open | Eyes Closed - Eyes Open | T | p |
|---|---|---|---|---|---|
| **Smoothing** | 0.59 | 0.35 | 0.24 | 17.18 | 2.2e-58 |
| **Smoothing+** | 0.49 | 0.30 | 0.19 | 16.20 | 8.2e-53 |
| **Basic** | 0.45 | 0.30 | 0.15 | 14.32 | 1.1e-42 |
| **Basic+** | 0.40 | 0.27 | 0.13 | 14.25 | 2.4e-42 |
| **CompCor** | 0.32 | 0.24 | 0.08 | 10.22 | 1.2e-23 |
| **CompCor+** | 0.29 | 0.22 | 0.07 | 10.45 | 1.2e-24 |

Supplementary Table 1. Significant differences in $GS_{amplitude}$ between segment types (EC > EO) for all pre-processing pipelines.

### Supplementary Methods

#### Critical Velocity

Based on formulation previously described by Kim et al. (2012), and used by Fultz et al. (2019), we compute the critical velocity of an inflow signal for a given slice as:

$$CV_s = 1000 \times \frac{d_s}{\Delta t_s}$$

where $d_s$ is the distance of slice $s$ to the lower edge of the imaging field of view in mm, and $\Delta t_s$ is the temporal gap between the start of the volume acquisition and the acquisition of slice $s$ in milliseconds.
